# Puberty contributes to adolescent development of fronto-striatal functional connectivity supporting inhibitory control

**DOI:** 10.1101/2022.05.02.490303

**Authors:** Amar Ojha, Ashley C. Parr, Will Foran, Finnegan J. Calabro, Beatriz Luna

## Abstract

Adolescence is defined by puberty and represents a period characterized by neural circuitry maturation (e.g., fronto-striatal systems) facilitating cognitive improvements. Though studies have characterized age-related changes, the extent to which puberty influences maturation of fronto-striatal networks is less known. Here, we combine two longitudinal datasets to characterize the role of puberty in the development of fronto-striatal resting-state functional connectivity (rsFC) and its relationship to inhibitory control in 106 10-18-year-olds. Beyond age effects, we found that puberty was related to decreases in rsFC between the caudate and the anterior vmPFC, rostral and ventral ACC, and v/dlPFC, as well as with rsFC increases between the dlPFC and nucleus accumbens (NAcc) across males and females. Stronger caudate rsFC with the dlPFC and vlPFC during early puberty was associated with worse inhibitory control and slower correct responses, respectively, whereas by late puberty, stronger vlPFC rsFC with the dorsal striatum was associated with faster correct responses. Taken together, our findings suggest that certain fronto-striatal connections are associated with pubertal maturation beyond age effects, which, in turn are related to inhibitory control. We discuss implications of puberty-related fronto-striatal maturation to further our understanding of pubertal effects related to adolescent cognitive and affective neurodevelopment.

## Introduction

Puberty demarcates the start of adolescence, the transitionary period of development to adulthood characterized by widespread biological, cognitive, and behavioral maturation. By the end of the adolescent period, individuals are better able to reliably engage brain circuitry that supports goal-directed cognitive processes (e.g., response inhibition) (Ordaz et al., 2013). In particular, inhibitory control continues to improve into the second decade of life as the percent of correct inhibitory responses increases and the latency to initiate a correct response decreases (Luna et al., 2004; Ordaz et al., 2017). Central to the development of these cognitive processes are maturational changes within fronto-striatal networks, comprised of prefrontal cortical (PFC) regions and striatal structures, such as the nucleus accumbens (NAcc), and their functional connectivity (Parr et al., 2021). What remains unclear, however, is the extent to which pubertal maturation contributes to the development of fronto-striatal circuitry to the adolescent transition to adult-level cognitive control beyond well-characterized age-related effects (Bos et al., 2012; van Duijvenvoorde et al., 2019; Fareri et al., 2015; Harsay et al., 2011; Parr et al., 2021). Studies in adolescents have largely relied on chronological age as the developmental variable of interest; however, investigating the effects of puberty (independent of age) represents an understudied area that is crucial to developing a more comprehensive understanding of the distinct mechanisms underlying adolescent neurodevelopment. Observations suggest that puberty has high inter-individual variability (Short and Rosenthal, 2008), is associated with psychopathological risks (Kuhn et al., 2010; Pfeifer and Allen, 2021), and posits specific biological mechanisms (e.g., sex hormones) underlying both the developmental improvements in behavior as well as increased risk for the emergence of psychopathology (e.g., mood disorders, substance use disorders, psychosis) (Paus et al., 2008) as well as sex differences in rates of mood and anxiety disorders during this period (Angold and Worthman, 1993). Pubertal maturation is strongly associated with psychopathological risk (Ho et al., 2021; Ladouceur et al., 2012; Mendle et al., 2010), particularly in girls (Oldehinkel et al., 2010), and some work suggests that puberty may better predict adolescent psychopathologies, such as substance use and depressive disorders, over and above age-related risks (Kuhn et al., 2010; Pfeifer and Allen, 2021). As such, investigating the unique contributions of puberty to neurodevelopment beyond age-related effects may provide a set of biological mechanisms that can clarify inter-individual differences in cognitive maturation across typical and atypical adolescent neurodevelopment.

Puberty—via its hormonal effects—impacts both physical maturation—resulting in the development of secondary sexual characteristics—and reproductive behaviors (Sisk and Foster, 2004). Importantly, it also contributes to cognitive changes, including in attentional and motivational processes (Hebbard et al., 2003; Romeo and Sisk, 2001; Sato et al., 2008). Animal studies have demonstrated that pubertal hormones are necessary for the formation of adult-like behavior. Interference with gonadal steroids during adolescence, via castration or pharmacological blockage, for example, produces deficits in adult-typical behaviors, (e.g., social interactions), even if hormones are later restored to typical levels in adulthood (Primus and Kellogg, 1990, 1989). Several studies have proposed that pubertal contributions to adolescent neurodevelopment occur, in part, via the influence of sex hormones on estrogen and estradiol receptors in the brain (Ho et al., 2020; Poon et al., 2019), which may mediate experience-dependent plasticity, allowing large-scale refinement and specialization of cognitive-affective networks, several of which may be especially sensitive to the stimulating and novel environments adolescents explore and exploit (Murty et al., 2016). These findings suggest that adolescence may reflect a critical period during which the effects of hormones on relevant neural circuitry determine cognitive, affective, and behavioral development (Ladouceur, 2012; Ladouceur et al., 2019; Larsen and Luna, 2018).

Central to facilitating goal-directed behaviors is the NAcc, which is innervated by dense dopaminergic projections originating in the ventral tegmental area (VTA) (Voorn et al., 1986). The NAcc undergoes substantial structural and functional maturation during adolescence (Larsen and Luna, 2014; Walhovd et al., 2014), and several human neuroimaging studies have observed increased NAcc activation during adolescence, potentially contributing to developmental “peaks” in reward-driven and risk-taking behaviors (Braams et al., 2015; Ernst et al., 2005; Galvan et al., 2006; Geier et al., 2009; Geier and Luna, 2012; Padmanabhan et al., 2011; Silverman et al., 2015). Further contributing to enhanced reward sensitivity, studies have shown heightened VTA – NAcc (Murty et al., 2018) and ventromedial prefrontal cortex (vmPFC) – NAcc (van Duijvenvoorde et al., 2019; Fareri et al., 2015; Parr et al., 2021) functional connectivity in adolescence during rewarded states that decreases into adulthood, paralleling developmental decreases in risk-taking behavior (Parr et al., 2021). Further, pubertal hormones such as testosterone, have been shown to mediate age-related changes in resting-state functional connectivity (rsFC) between the NAcc and medial PFC (Fareri et al., 2015), suggesting that NAcc circuitry bridging midbrain dopamine systems and prefrontal decision-making and/or control regions may be particularly sensitive to pubertal maturational processes. Along with the NAcc, the dorsal striatum—and the caudate nucleus, in particular—has been implicated in response inhibition (Harsay et al., 2011; Zhang et al., 2017), undergoes developmental changes in functional connectivity during adolescence (van Duijvenvoorde et al., 2019), and has been observed to be associated with psychopathology, such as with adolescent suicidality (Harms et al., 2019; Ho et al., 2021); however, the majority of the human neuroscience studies in adolescents have not considered the extent to which puberty— above and beyond age-related effects—relates to the development of neural systems implicated in cognitive maturation, such as fronto-striatal circuitry.

The present study seeks to bridge this gap in the field by characterizing and understanding the role of puberty in fronto-striatal contributions to cognitive development. Using neuroimaging and behavioral data from two longitudinal adolescent samples, we investigated the effects of pubertal maturation on fronto-striatal rsFC after accounting for age-related effects. Based on previous developmental findings, we hypothesized that pubertal maturation would be associated with rsFC decreases between medial frontal areas (e.g., ventromedial PFC (vmPFC) and anterior cingulate cortex (ACC) subregions) and the NAcc (Parr et al., 2021) and the caudate (van Duijvenvoorde et al., 2019), which have also been implicated in reward sensitivity (Blakemore, 2008; Crone, 2014; van Duijvenvoorde et al., 2015; Jones et al., 2014). In contrast, we expect lateral frontal areas (e.g., dorsolateral and ventrolateral PFC (dlPFC, vlPFC)) rsFC with the NAcc and caudate to increase with pubertal maturation, given their roles in improvements related to executive functions, such as cognitive control (MacDonald et al., 2000), response inhibition (Luna et al., 2001), and age-related decreases in risk-taking behaviors (Qu et al., 2015). To test the specificity of puberty-related associations with maturation of striatal connectivity, we additionally investigate the putamen but do not expect to observe puberty-related changes in putamen connections. Furthermore, we expect that female participants will exhibit earlier puberty-related fronto-striatal functional maturation relative to male participants, given their relatively earlier timing of pubertal development. Finally, given evidence implicating these networks in developmental changes in cognition (Morein-Zamir and Robbins, 2014) through adolescence, we will explore the association between puberty-related maturation within fronto-striatal networks and inhibitory control performance as measured by the antisaccade task. Taken together, this work will contribute to deepening our understanding of the role of puberty as it relates to cognitive improvements observed during the adolescent period and may inform some of the sex differences observed in the incidence rates for affective psychopathologies during this critical period of development (Dalsgaard et al., 2020).

## Methods

### 2.1. Participants

Participants were drawn from two large, longitudinal neuroimaging studies, which have been previously reported (Calabro et al., 2019; Parr et al., 2022, 2021). For our primary analyses, 106 adolescents (50 females) ages 10-18 provided 198 usable resting-state scan data and reported starting puberty over 1-3 longitudinal visits (n = 50 provided data from one visit) at 12-18 month intervals (details below). One of the cohorts was also examined in a separate paper investigating puberty effects on wide-ranging connectivity and inhibitory control performance (Ravindranath et al., 2022, submitted to this issue). Participants were recruited from the community and screened for the absence of neurological or psychiatric problems including loss of consciousness, self or first-degree relatives with major psychiatric illness, and MRI scanning contraindications (e.g., claustrophobia, metal in body, pregnancy). Participants under the age of 18 provided assent and parents of participants provided informed consent. Experimental procedures were approved by the University of Pittsburgh Institutional Review Board and complied with the Declaration of Helsinki. Participants were compensated for their time.

To contextualize findings across adolescence and into early adulthood, we also characterized age effects in participants under age 18, including in those without pubertal data (161 participants, 87 females; 289 scans), as well as across all participants (up to age 35) who provided usable rsfMRI data and completed up to 13 longitudinal visits (285 participants, 153 females; 763 scans).

### 2.2. Puberty assessment

Pubertal assessments were collected in participants ages 18 and under using the Petersen Pubertal Development Scale (PDS) (Petersen et al., 1988), a self-report measure of physical development. The PDS is comprised of five questions about physical development, differing for males and females, with possible item scores ranging from 1 (development has not yet begun) to 4 (development appears complete), and includes several additional (secondary) sexual characteristic questions not captured by Tanner staging (e.g., skin and hair changes). Composite scores were transformed to a 5-point scale for comparison to Tanner scale stages. Given our interest in the role of pubertal maturation, participants were included in the puberty analyses if their transformed-PDS score indicated pubertal development was already underway (transformed-PDS score ≥ 2). See **Fig. 1** for participant distributions by age, puberty, and sex. Given the skewness of our sample’s pubertal data, we considered log-transformations of puberty; however, given that this normalization approach using log transformations did not reduce skewness in our sample, we did not further transform the distribution of puberty data.

**Figure 1.**
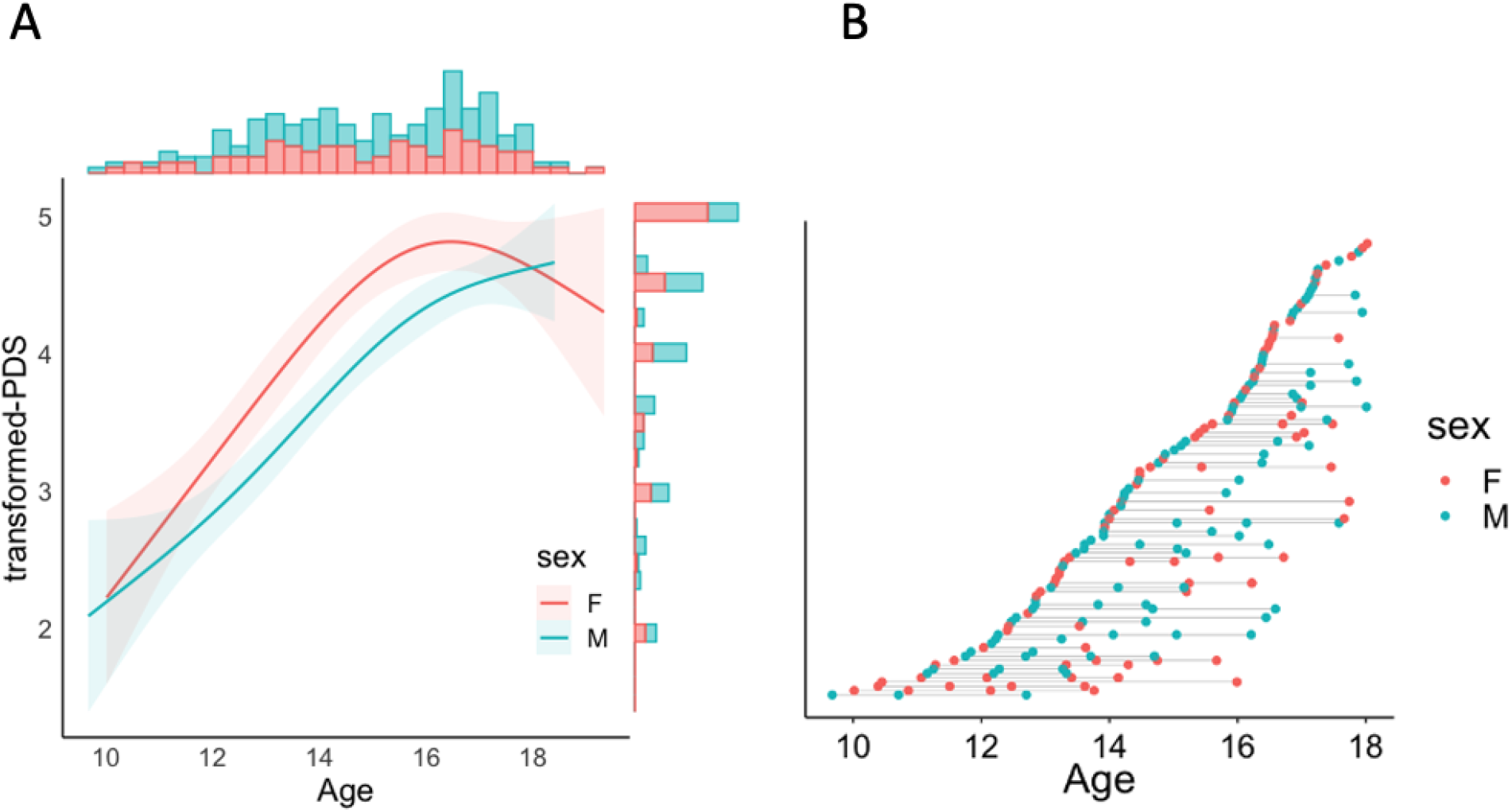
Distribution of participants by (**A**) age and pubertal maturation (transformed-PDS), separated by sex, along with (**B**) a plot depicting the number of visits per participant: each point represents an individual scan, with lines connecting individual participants.

### 2.3 Eye tracking data acquisition

Eye tracking data were captured using an eye-tracking system (M5000, Applied Science Laboratories) with a sampling rate of 60 Hz. Real-time monitoring enabled identification of head movement or inattention to the task, in which case, experimenters redirected subjects following the run. A 9-point calibration routine was performed at the beginning of each session. Responses were scored with a custom scoring script written in R (see Ravindranath et al., 2020 for details).

### 2.4 Antisaccade task

Participants completed a total of 48 antisaccade (AS) trials at each visit, performed outside of the scanner (previously described in Ordaz et al., 2013 and Ravindranath et al., 2020). Each AS trial began with a red fixation cross presented at the center of a black screen for 500-6000 ms followed by a 200 ms black screen. Next, a yellow “cue” dot appeared pseudo-randomly (evenly distributed between four positions on the horizonal meridian at 2%, 33%, 66%, or 98% of the screen width) on the screen for 1000 ms along the horizontal axis in the center of the screen followed, by a 20 ms response period (black screen) during which participants were instructed to direct their gaze away from the location of the yellow cue and instead, look at the mirror location on the screen. See **Fig. 2** for a schematic of the antisaccade task.

**Figure 2.**
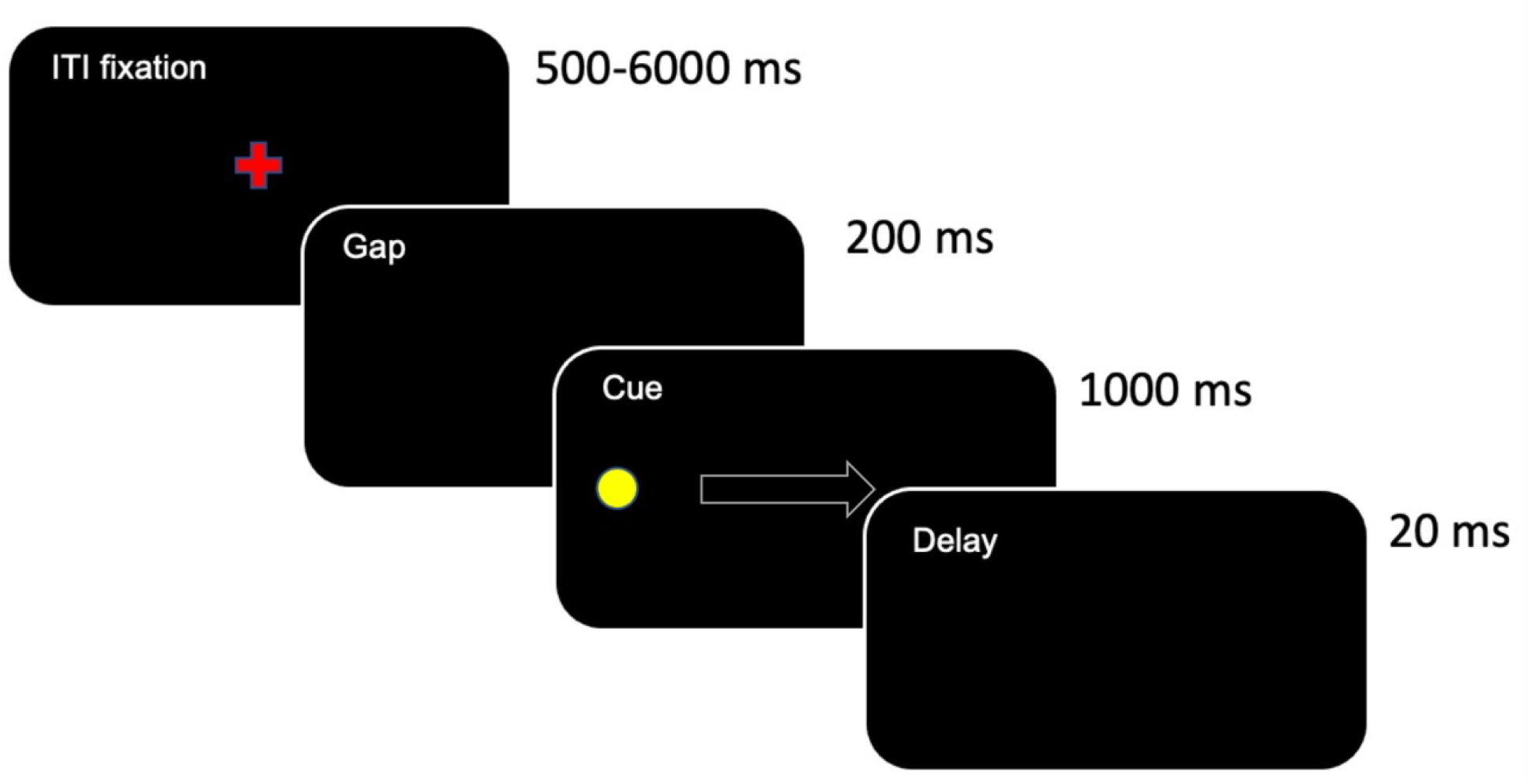
Antisaccade task schematic.

AS responses were considered “correct” if the first eye movement during the saccade epoch had a velocity ≥ 30°/second toward the mirror location of the yellow cue and extended beyond a 2.5°/visual angle from central fixation. A trial was considered an error if the first saccade in the response epoch was directed toward the peripheral stimulus and extended beyond 2.5° central fixation window. Trials were considered error-corrected if an error was followed by a correct response. Express saccades, which are too rapid to engage cognitive systems, were excluded and defined as a saccade starting within the first 4 samples (60Hz) after trial onset. For the current study, we focused our analyses on correct antisaccade trials to understand pubertal and neural associations with adult-like cognitive maturation, as evidenced by successful inhibitory response execution (i.e., making a correct antisaccade eye movement), which has been shown to improve through adolescence and stabilize in adulthood (Luna et al., 2004; Murty et al., 2018; Ordaz et al., 2013; Simmonds et al., 2017). Our two inhibitory control variables of interest were antisaccade “performance” (number of correct / total usable trials) and mean latency—the time from when the stimulus appeared to saccade onset (in ms)—on correct trials.

### 2.5. MR data acquisition

#### MMR dataset

MR data were acquired on a 3T Siemens Biograph molecular Magnetic Resonance (mMR) PET/MRI scanner. Structural images were acquired using a T1 weighted magnetization-prepared rapid gradient-echo (MPRAGE) sequence (TR, 2300 ms; echo time (TE), 2.98 ms; flip angle, 9°; inversion time (TI), 900 ms, voxel size, 1.0 × 1.0 × 1.0 mm). Functional images were acquired using blood oxygen level dependent (BOLD) signal from an echoplanar sequence (TR, 1500 ms; flip angle, 50°; voxel size, 2.3 × 2.3 × 2.3 mm in-plane resolution) with contiguous 2.3mm – thick slices aligned to maximally cover the cortex and basal ganglia. Two 8-min sessions of fixation resting-state fMRI (rsfMRI) data were collected prior to and following a task-based fMRI sequence, respectively.

#### CogLong dataset

MR data were acquired on a Siemens 3T MAGNETOM Allegra. Structural images were acquired using a T1 weighted magnetization-prepared rapid gradient-echo (MPRAGE) sequence (TR, 1570 ms; TE, 3.04 ms; flip angle, 8°; TI, 800 ms, voxel size, 0.78125 × 0.78125 × 1.0 mm). Functional images were acquired using BOLD signal from an echoplanar sequence (TR, 1500 ms; TE, 25 ms; flip angle, 70°; voxel size, 3.125 × 3.125 mm in-plane resolution) with contiguous 4-mm-thick slices aligned to the subject’s anterior-posterior commissure plane.

rsfMRI data were extracted from the OFF periods of a mixed block-event-related design (Ordaz et al., 2013) based on a previously reported method (Fair et al., 2007), which found that fMRI data from blocked design tasks with rest periods are largely similar to rsfMRI data collected continuously. Briefly, during ON periods, participants completed either a pro- or anti-saccade task. Functional images were extracted from the OFF periods, excluding the 15s following the preceding ON period. Across the four runs, 6min48s min of rsfMRI data were produced, thereby providing enough time for signal stabilization necessary for resting-state data analysis (Dijk et al., 2010). See (Calabro et al., 2019) for additional details.

### 2.6. MR data preprocessing

Structural MRI data were preprocessed to extract the brain from the skull and warped to the MNI standard using both linear (FLIRT) and non-linear (FNIRT) transformations. rsfMRI data were preprocessed using a pipeline that minimized the effects of head motion (Hallquist et al., 2013) including 4D slice-timing and head motion correction, skull stripping, intensity thresholding, wavelet despiking (Patel et al., 2014), coregistration to the structural image and nonlinear warping to MNI space, local spatial smoothing with a 5mm Gaussian kernel based on the SUSAN algorithm, intensity normalization, and nuisance regression based on head motion (6° of translation/rotation and their first derivative) and non-gray matter signal (white matter and CSF and their first derivative). Bandpass filtering between .009 and .08 Hz was done simultaneously with nuisance regression. Frame-wise motion estimates were computed for resting-state data. Functional volumes containing frame-wise displacement (FD) > 0.3 mm were excluded from analyses (Siegel et al., 2013). Participants with more than 40% of TRs censored were excluded altogether from rsfMRI analyses, resulting in the exclusion of 64 participants. Neuroimaging analyses were performed in AFNI (Cox, 1996).

### 2.7. Region of interest (ROI) selection

We used a region of interest (ROI) approach to investigate fronto-striatal connectivity between hypothesis-driven, *a priori* ROIs. Striatal ROIs included the NAcc, a central node of reward networks previously shown to support goal-directed behavior (Mannella et al., 2013; Parr et al., 2021), including performance on the antisaccade task (Murty et al., 2018), as well as the caudate nucleus, given previous reports demonstrating age-related maturation of this region to be associated with aspects of response inhibition and cognitive control (Hu et al., 2018; Jahfari et al., 2011; Rubia et al., 2006). Although we were mainly interested in the NAcc and caudate as striatal ROIs, we also examined the putamen as an additional striatal region to test the specificity of pubertal effects on fronto-striatal connections more broadly.

We selected frontal regions known to support cognitive and affective maturation during adolescence, including the dlPFC, vlPFC, anterior vmPFC, and subgenual, rostral, and ventral ACC (sgACC, rACC, vACC, respectively) (Ernst et al., 2005; Galván et al., 2006; Van Leijenhorst et al., 2010; Geier et al., 2010; Padmanabhan et al., 2011; Parr et al., 2011). Medial structures, which included the anterior vmPFC, sgACC, rACC, and vACC, were defined based on the Mackey and Petrides atlas (Mackey and Petrides, 2014). We excluded orbitofrontal cortex (OFC) ROIs from analyses due to relatively lower temporal signal-to-noise ratios (tSNRs) for these PFC subregions relative to the other ROIs despite their contributions to goal-directed behaviors (Rudebeck and Rich, 2018). The dlPFC ROI was defined using the MNI Glasser Human Connectome Project atlas (Glasser et al., 2016). The vlPFC and all striatal ROIs were defined using the Brainnetome atlas (Fan et al., 2016). Voxels needed to have at least a 50% or greater probability of being gray matter in the MNI-152-09c template to be included in each ROI. The final frontal ROIs included the anterior vmPFC (area 14m), sgACC (area 25), rACC (area 32), vACC (area 24), dlPFC (areas 8, 9, and 46), and vlPFC (areas 44, 45, and 47). See **Fig. 3** for a representation of these ROIs.

**Figure 3.**
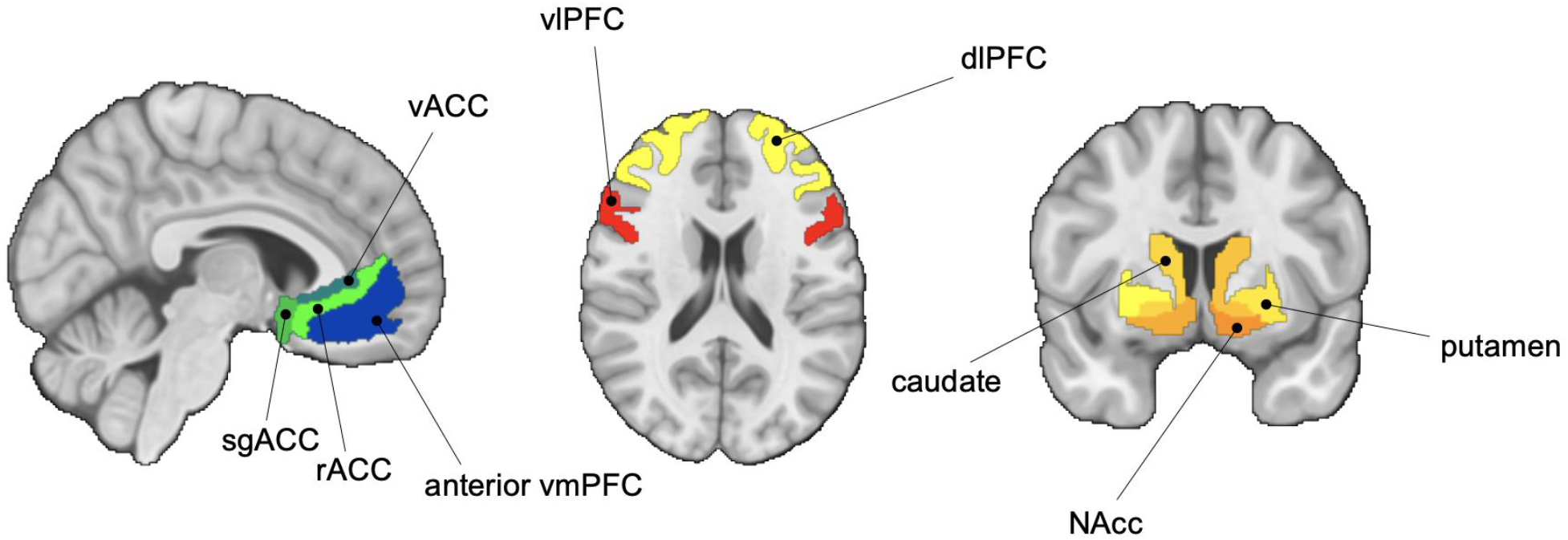
Fronto-striatal regions of interest (ROIs). Abbreviations: NAcc, nucleus accumbens; vmPFC, ventromedial prefrontal cortex; sgACC, subgenual cingulate; vACC, ventral anterior cingulate cortex; rACC, rostral anterior cingulate cortex; dlPFC, dorsolateral prefrontal cortex; vlPFC, ventrolateral prefrontal cortex.

### 2.8. Functional connectivity analyses

#### ROI-based

Time series were extracted from each participant’s preprocessed resting-state functional images by taking the first principal component across all voxels within each ROI (Zhou et al., 2009). Pearson correlation coefficients were then computed between ROI seeds and normalized using Fisher’s Z transformation. If both of a participant’s rsfMRI scans from the MMR dataset were usable, we averaged rsFC values for each connectivity pair. In cases where only one scan was usable, the unusable scan was dropped from analyses. rsFC data were harmonized between the two datasets (i.e., MMR, CogLong) using NeuroComBat (*neuroCombat* in R, R Core Team, 2020) to account for scanner acquisition differences while retaining biological variability of interest (Fortin et al., 2018).

#### Seed-based

To ensure that our results were not biased based on analytic choices (e.g., frontal ROI definition), we additionally performed exploratory seed-based rsFC analyses using AFNI’s *3dLMEr, 3dFWHMx*, and *3dClustSim*. As these analyses were exploratory, we examined each of the three striatal ROIs by hemisphere to detect any laterality effects we may have missed using our bilateral ROI approach.

### 2.9 Statistical analysis

We used generalized additive mixed models (GAMMs; *gamm4* package; R version 4.0.3 via RStudio version 2021.09.1; R Core Team, 2020) for fronto-striatal developmental (i.e., age, puberty) analyses, including a random offset per participant to account for longitudinal repeated measures. GAMMs are similar to generalized linear mixed models but also include smooth functions for predictor variables (“smooth splines”), which are sensitive to linear and non-linear effects, but penalize for additional parameters to prevent overfitting models (Wood, 2013; Wood et al., 2017). We ran GAMMs testing the main effects of puberty on fronto-striatal rsFC and modeled age and sex as covariates (Model 1 below). We controlled for laterality and hemisphere effects by including one term for frontal ROI hemisphere (either ‘right’ or ‘left’), as well as a laterality term for the connectivity pair (either ‘ipsilateral’ or ‘contralateral’)—similar to past approaches we have used to leverage these data to increase statistical power (Parr et al., 2021; Tervo-Clemmens et al., 2017) —given our lack of hypotheses regarding lateralization effects. As such, our primary analyses were modeled as follows:

Model 1:

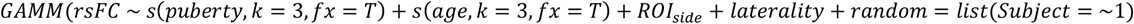

We additionally tested for puberty-by-sex interactions (Model 2 below) and, in cases of significant findings, followed up with post-hoc tests investigating puberty effects by sex in males and females, and again controlled for age, hemisphere, and laterality effects.

Model 2:

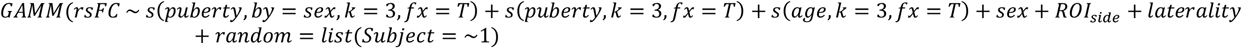

We used a similar approach to characterize age effects (without the puberty term) in participants up to age 18, and then, separately, to characterize age effects up to age 35. All reported *p*-values are Bonferroni corrected to account for multiple comparisons. Finally, to test whether associations between fronto-striatal rsFC and AS performance and latency on correct trials differed as a function of puberty, we used linear mixed effects models (*lme4* package in R) to test for interaction effects in the significant puberty-related connections. If we had observed a significant puberty-by-sex interaction on fronto-striatal rsFC, we tested the associations between rsFC and AS as a function of puberty separately in males and females. In cases in which we only observed a significant main effect of puberty, we modeled sex as a covariate term and tested relationships across males and females. We again used the hemisphere and laterality covariate approach described above and included age^−1^ as an additional covariate, given previous findings suggesting age-related improvements on the AS are characterized by inverse functions (Luna, 2009; Luna et al., 2004).

## Results

### 3.1. Puberty-related fronto-striatal functional connectivity effects

#### 3.1.1. Puberty-related effects on fronto-striatal rsFC

We characterized the effects of age and puberty on fronto-striatal rsFC by testing for age effects in both the full (up to age 35) and pubertally developing (up to age 18) samples (see **Supplemental Results** for age effects), and by testing both main effects of puberty and puberty by sex interactions after controlling for these age effects.

##### Main effects of puberty

We found several connections that exhibited main effects of puberty (**Fig. 4, Fig. S1**, and **Table 1**). Across males and females, we observed a significant main effect of puberty beyond age-related effects for NAcc connectivity with the vlPFC (*F* = 11.39 *p*_Bonferroni_ = .0002) and dlPFC (*F* = 13.59, *p*_Bonferroni_ < .0001). We also found significant puberty effects between the caudate and vlPFC rsFC (*F* = 10.02, *p*_Bonferroni_ = .0009), anterior vmPFC (*F* = 7.04, *p*_Bonferroni_ = .017), rACC (*F* = 7.58, *p*_Bonferroni_ = .010), and vACC (*F* = 10.11, *p*_Bonferroni_ = .0008). Finally, the putamen also showed a significant puberty-related decrease in its connectivity with vlPFC (*F* = 6.90, *p*_Bonferroni_ = .019). All of the significant connections described above exhibited puberty-related decreases in rsFC from approximately stages 2 – 3.5, followed by either a plateau, or, as in the case of dlPFC – NAcc, an increase in rsFC from approximately stages 3.5 – 5. No other significant main effects of puberty were observed in any other fronto-striatal connection when controlling for age and sex effects (*p*s > .05). See **Supplemental Results** for seed-based analyses and sensitivity analyses covarying for motion (**Table S1**). Briefly, exploratory striatal seed-based analyses revealed no additional frontal clusters associated with clusters. Further, no significant relationship changed after controlling for motion (percent of TRs censored).

**Figure 4.**
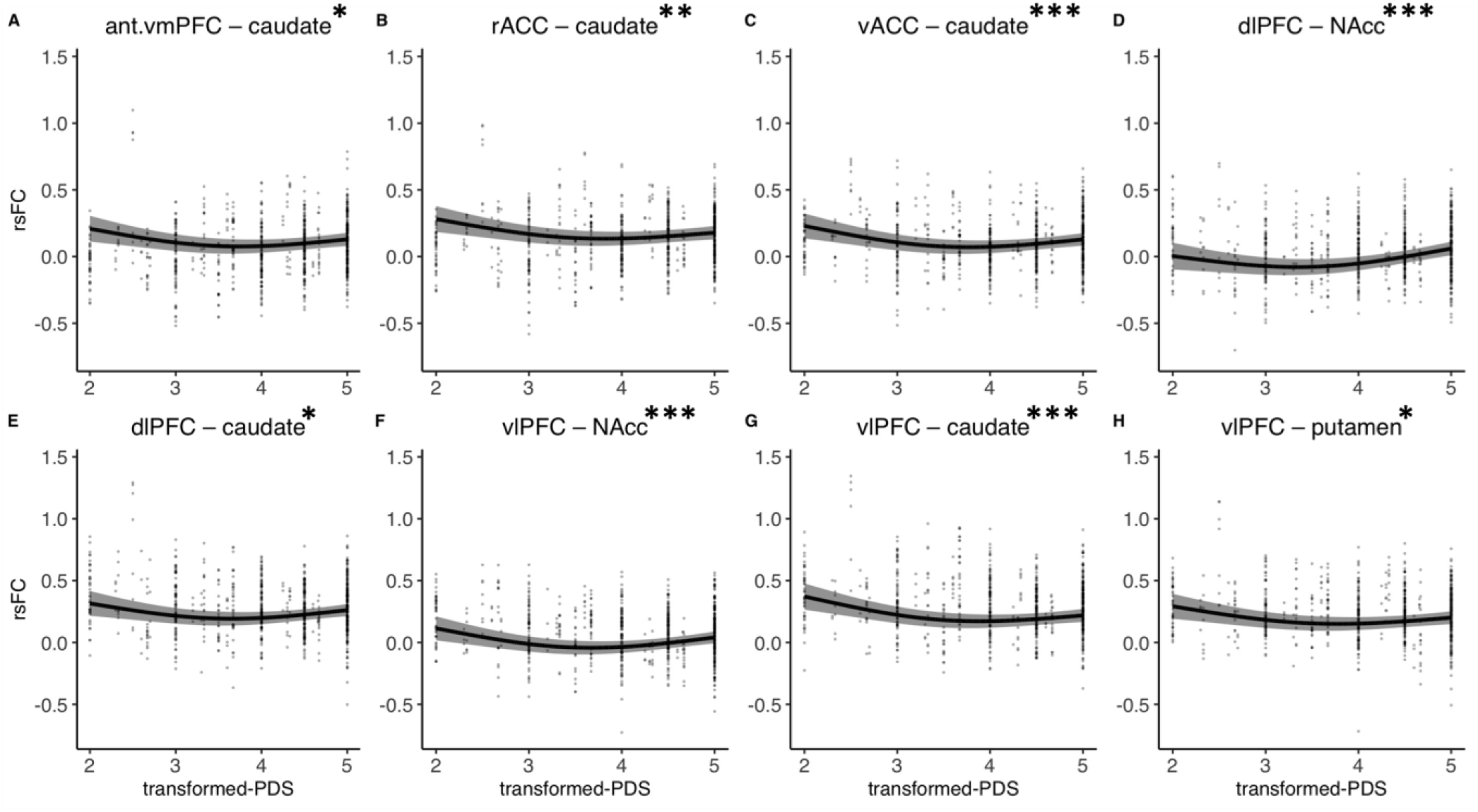
Main effects of puberty on fronto-striatal resting-state functional connectivity (rsFC) beyond age and sex effects. Abbreviations: NAcc, nucleus accumbens; vmPFC, ventromedial prefrontal cortex; sgACC, subgenual cingulate; vACC, ventral anterior cingulate cortex; rACC, rostral anterior cingulate cortex; dlPFC, dorsolateral prefrontal cortex; vlPFC, ventrolateral prefrontal cortex; PDS, Petersen Pubertal Development Scale. * *p*_Bonferroni_ < .05, ** *p*_Bonferroni_ < .01, *** *p*_Bonferroni_ < .001.

**Table 1.**
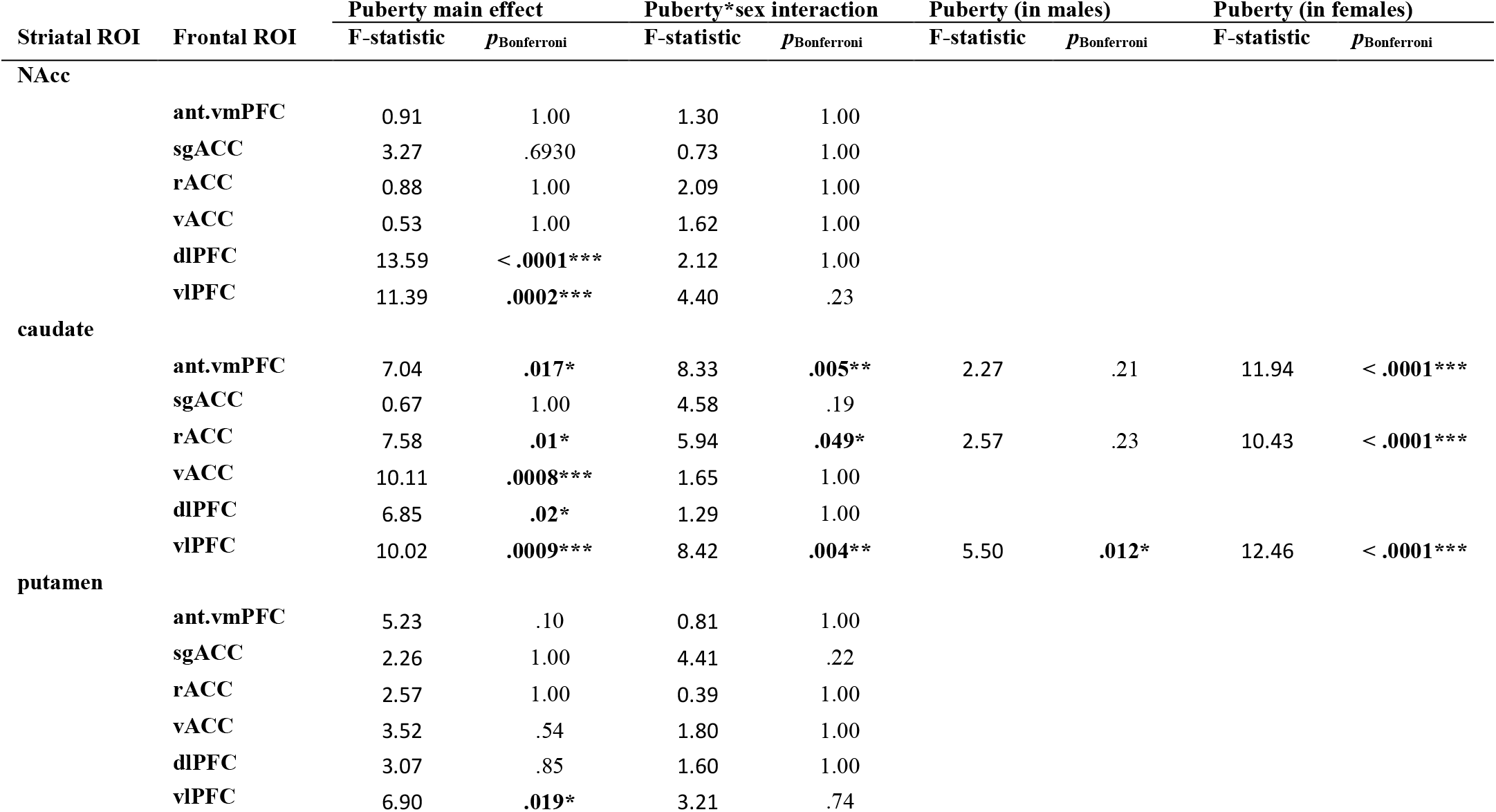
Generalized Additive Mixed-effects Models (GAMMs) examining associations between puberty and fronto-striatal resting-state functional connectivity. Age and sex were modeled as covariates in main and interaction models. Post-hoc within sex tests were performed only for fronto-striatal connections exhibiting significant puberty-by-sex interactions. * *p*_Bonferroni_ < .05, ** *p*_Bonferroni_ < .01, *** *p*_Bonferroni_ < .001 Abbreviations: ant. vmPFC, anterior ventromedial prefrontal cortex; sgACC, subgenual anterior cingulate cortex; rACC, rostral anterior cingulate cortex; vACC, ventral anterior cingulate cortex; dlPFC, dorsolateral prefrontal cortex; vlPFC, ventrolateral prefrontal cortex.

##### Puberty-by-sex interaction effects

We observed significant puberty-by-sex interactions (beyond age-related effects) in three fronto-caudate connections (**Fig. 5, Fig. S2**, and **Table 1**), including connectivity between the caudate and the anterior vmPFC (*F* = 8.33, *p*_Bonferroni_ = .005), rACC (*F* = 5.94, *p*_Bonferroni_ = .049), and vlPFC (*F* = 8.42, *p*_Bonferroni_ = .004). Post-hoc tests examining puberty effects within each sex for these connections revealed a significant puberty effect in females for caudate rsFC with the anterior vmPFC (*F* = 11.94, *p*_Bonferroni_ < .0001), rACC (*F* = 10.43, *p*_Bonferroni_ = .0001), and vlPFC (*F* = 12.46, *p*_Bonferroni_ < .0001). Caudate rsFC with both the anterior vmPFC and rACC was characterized by increases in connectivity strength in late puberty (approximately stages 4 – 5), while vlPFC – caudate rsFC in females was characterized by a quadratic (U-shaped) effect, exhibiting puberty-related decreases from stages 2 – 3.5, followed by increases in late puberty (stages 4 – 5). In contrast, we only observed one significant puberty effect in males of the three connections tested: puberty was associated with a linear shaped decrease in vlPFC – caudate rsFC (*F* = 5.50, *p*_Bonferroni_ = .012).

**Figure 5.**
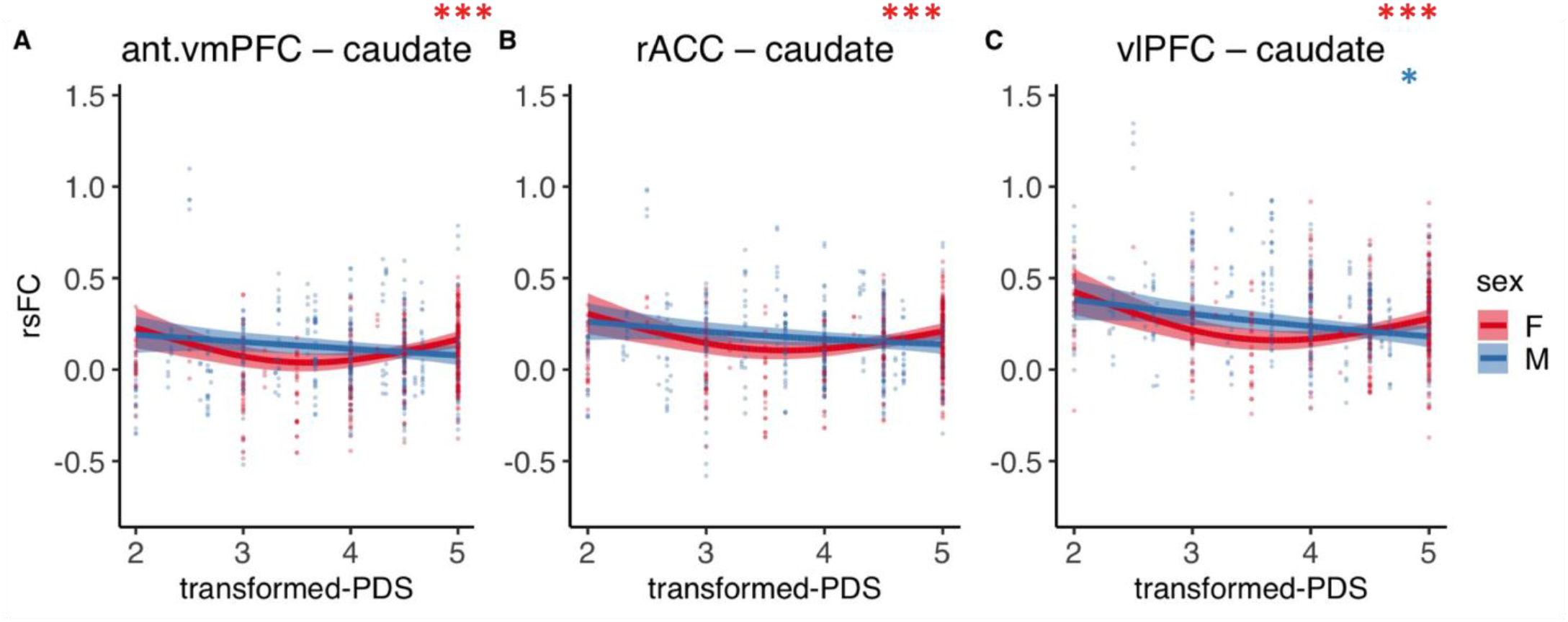
Puberty-by-sex interaction effects beyond age on fronto-striatal resting-state functional connectivity (rsFC). Asterisks denoting significance refer to post-hoc within sex puberty effects. Abbreviations: ant.vmPFC, anterior ventromedial prefrontal cortex; rACC, rostral anterior cingulate cortex; vlPFC, ventrolateral prefrontal cortex; PDS, Petersen Pubertal Developmental Scale; rsFC, resting-state functional connectivity. * *p*_Bonferroni_ < .05, ** *p*_Bonferroni_ < .01, *** *p*_Bonferroni_ < .001.

### 3.2. Age-related fronto-striatal functional connectivity effects

Main effects of age as well as age-by-sex interaction effects on fronto-striatal rsFC are presented in **Tables 2** and **3** for the pubertal (<18yo) and full (<35yo) samples, respectively. Plots illustrating age effects are presented in **Fig. S2** and **S3** for the pubertal (<18yo) sample for main effects of age and age-by-sex interaction effects, respectively. Plots illustrating age effects are presented in **Fig. S4** and **S5** for the full (<35yo) sample for main effects of age and age-by-sex interaction effects, respectively.

**Table 2.** Generalized Additive Mixed-effects Models (GAMMs) examining associations between age (up to 18-years-old) and fronto-striatal resting-state functional connectivity. Sex was modeled as covariates in main and interaction models. Post-hoc within sex tests were performed only for fronto-striatal connections exhibiting significant age-by-sex interactions. *p*_Bonferroni_ < .05, ** *p*_Bonferroni_ < .01, *** *p*_Bonferroni_ < .001

| Striatal ROI | Frontal ROI | Age main effect |  | Age*sex interaction |  | Age (in males) |  | Age (in females) |  |
| --- | --- | --- | --- | --- | --- | --- | --- | --- | --- |
|  |  | F-statistic | *p*<sub>Bonferroni</sub> | F-statistic | *p*<sub>Bonferroni</sub> | F-statistic | *p*<sub>Bonferroni</sub> | F-statistic | *p*<sub>Bonferroni</sub> |
| NAcc |  |  |  |  |  |  |  |  |  |
|  | ant.vmpFC | 7.10 | **0.016\*** | 1.27 | 1.00 |  |  |  |  |
|  | sgACC | 2.95 | 0.95 | 0.97 | 1.00 |  |  |  |  |
|  | rACC | 3.62 | 0.49 | 0.74 | 1.00 |  |  |  |  |
|  | vACC | 4.89 | 0.14 | 0.45 | 1.00 |  |  |  |  |
|  | dIPFC | 1.60 | 1.00 | 0.23 | 1.00 |  |  |  |  |
|  | vIPFC | 5.46 | 0.079 | 1.97 | 1.00 |  |  |  |  |
| caudate |  |  |  |  |  |  |  |  |  |
|  | ant.vmpFC | 6.58 | **0.026\*** | 7.87 | **0.007\*\*** | 13.35 | **< .0001\*\*\*** | 0.45 | 1.00 |
|  | sgACC | 0.61 | 1.00 | 3.12 | 0.80 |  |  |  |  |
|  | rACC | 6.65 | **0.024\*** | 24.89 | **< .0001\*\*\*** | 29.41 | **< .0001\*\*\*** | 1.66 | 1.00 |
|  | vACC | 0.45 | 1.00 | 11.34 | **0.0002\*\*\*** | 7.43 | **0.004\*\*** | 4.32 | 0.08 |
|  | dIPFC | 3.71 | 0.45 | 1.91 | 1.00 |  |  |  |  |
|  | vIPFC | 7.14 | **0.015\*** | 4.08 | 0.31 |  |  |  |  |
| putamen |  |  |  |  |  |  |  |  |  |
|  | ant.vmpFC | 0.87 | 1.00 | 0.002 | 1.00 |  |  |  |  |
|  | sgACC | 0.97 | 1.00 | 2.49 | 1.00 |  |  |  |  |
|  | rACC | 0.68 | 1.00 | 1.64 | 1.00 |  |  |  |  |
|  | vACC | 0.45 | 0.50 | 0.25 | 1.00 |  |  |  |  |
|  | dIPFC | 2.96 | 0.94 | 1.43 | 1.00 |  |  |  |  |
|  | vIPFC | 12.9 | **0.0001\*\*\*** | 3.73 | 0.44 |  |  |  |  |

**Table 3.** Generalized Additive Mixed-effects Models (GAMMs) examining associations between age (up to 35-years-old) and fronto-striatal resting-state functional connectivity. Sex was modeled as covariates in main and interaction models. Post-hoc within sex tests were performed only for fronto-striatal connections exhibiting significant age-by-sex interactions. *p*_Bonferroni_ < .05, ** *p*_Bonferroni_ < .01, *** *p*_Bonferroni_ < .001

| Striatal ROI | Frontal ROI | Age main effect |  | Age*sex interaction |  | Age (in males) |  | Age (in females) |  |
| --- | --- | --- | --- | --- | --- | --- | --- | --- | --- |
| | | F-statistic | $p_{\text{Bonferroni}}$ | F-statistic | $p_{\text{Bonferroni}}$ | F-statistic | $p_{\text{Bonferroni}}$ | F-statistic | $p_{\text{Bonferroni}}$ |
| NAcc | ant.vmpFC | 5.75 | 0.06 | 21.14 | < .0001*** | 10.47 | 0.0002*** | 16.59 | < .0001*** |
|  | sgACC | 16.77 | < .0001*** | 2.27 | 1.00 |  |  |  |  |
|  | rACC | 1.72 | 1.00 | 18.01 | < .0001*** | 6.63 | 0.008** | 13.18 | < .0001*** |
|  | vACC | 1.20 | 1.00 | 6.88 | 0.019* | 3.71 | .15 | 4.53 | 0.065 |
|  | dIPFC | 16.03 | < .0001*** | 3.40 | 0.61 |  |  |  |  |
|  | vIPFC | 0.62 | 1.00 | 0.98 | 1.00 |  |  |  |  |
| caudate | ant.vmpFC | 31.48 | < .0001*** | 22.24 | < .0001*** | 48.28 | < .0001*** | 6.07 | 0.012* |
|  | sgACC | 1.47 | 1.00 | 0.34 | 1.00 |  |  |  |  |
|  | rACC | 16.18 | < .0001*** | 11.04 | < .0001*** | 25.23 | < .0001*** | 2.31 | 0.59 |
|  | vACC | 3.95 | 0.35 | 2.63 | 1.00 |  |  |  |  |
|  | dIPFC | 1.78 | 1.00 | 1.87 | 1.00 |  |  |  |  |
|  | vIPFC | 3.65 | 0.47 | 3.84 | 0.39 |  |  |  |  |
| putamen | ant.vmpFC | 6.21 | 0.036* | 7.91 | 0.007** | 10.52 | 0.0002*** | 3.57 | 0.17 |
|  | sgACC | 1.83 | 1.00 | 0.21 | 1.00 |  |  |  |  |
|  | rACC | 3.07 | 0.84 | 2.53 | 1.00 |  |  |  |  |
|  | vACC | 4.55 | 0.19 | 0.20 | 1.00 |  |  |  |  |
|  | dIPFC | 17.54 | < .0001*** | 1.66 | 1.00 |  |  |  |  |
|  | vIPFC | 7.40 | 0.011* | 0.64 | 1.00 |  |  |  |  |

### 3.3. Developmental effects on antisaccade (AS) performance and latency

AS performance (i.e., proportion of correct trials) significantly improved with both age (pubertal <18yo sample: *β* = .50, *t* = 7.78, *p* < .0001, **Fig. S7**; full <35yo sample: *β* = .32, *t* = 7.19, *p* < .0001, **Fig. S8**) and pubertal stage (*β* = .33, *t* = 3.94, *p* = .0001, **Fig. 6**) after covarying for sex. The effect of pubertal stage was diminished after controlling for age (*β* = .02, *t* = .17, *p* = .86), consistent with previous findings (Ordaz et al., 2017). We did not observe a significant age-by-sex (pubertal <18yo sample: *β* = .12, *t* = .91, *p* = .37; full <35yo sample: *β* = .01, *t* = .16, *p* = .87) or puberty-by-sex (*β* = .22, *t* = 1.32, *p* = .19) interaction, including after controlling for age (*β* = .20, *t* = 1.25, *p* = .21), on AS performance.

**Figure 6.**
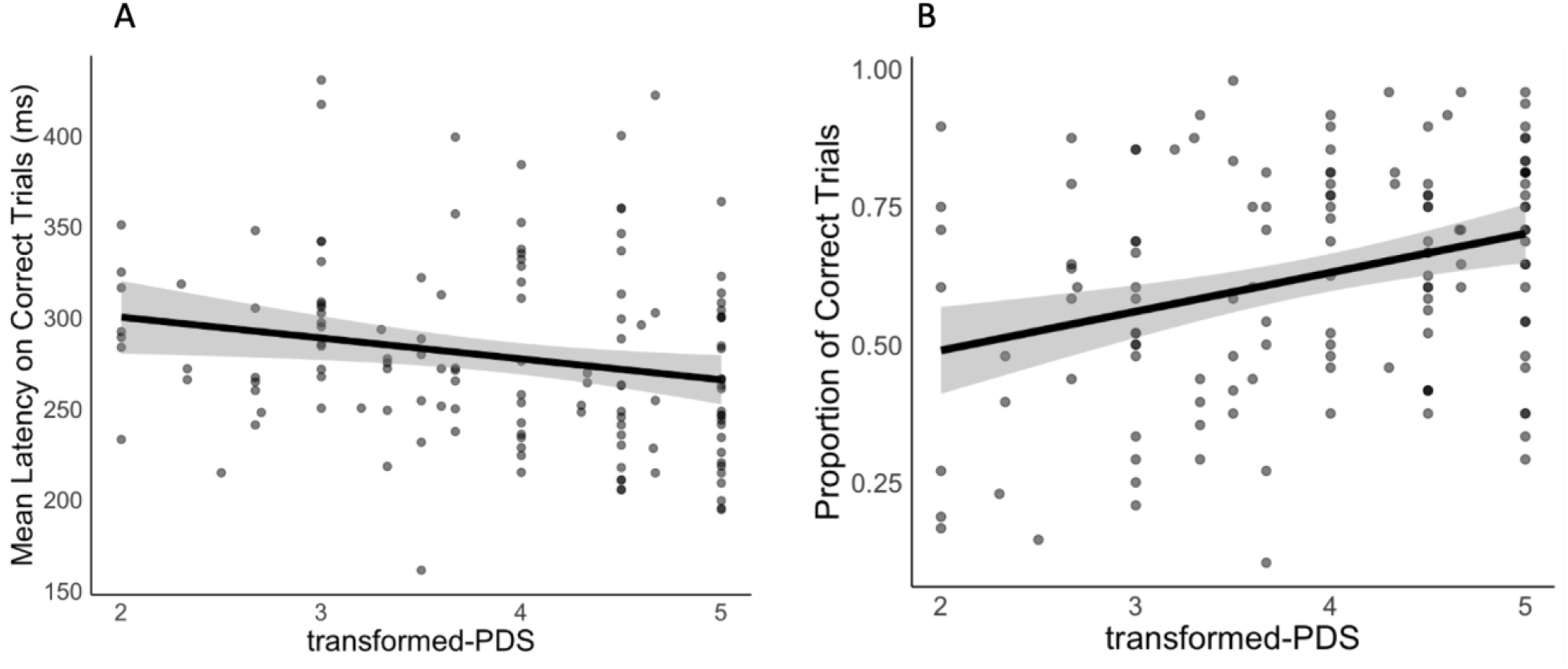
Antisaccade task performance on correct trials by pubertal maturation separated by (**A**) mean latency on correct trials (in ms) and (**B**) proportion of correct trials.

AS latency (on correct trials) significantly decreased with both age (pubertal <18yo sample: *β* = -.58, *t* = -8.68, *p* < .0001; full <35yo sample: *β* = -.48, *t* = -11.19, *p* < .0001) and pubertal stage (*β* = -.24, *t* = -2.51, *p* = .014) after controlling for sex, but again the effect of pubertal stage was diminished after controlling for age (*β* = .13, *t* = .90, *p* = .37). We did not observe a significant age-by-sex (pubertal <18yo sample: *β* = .04, *t* = .31, *p* = .76; full <35yo sample: *β* = .04, *t* = .51, *p* = .61) or puberty-by-sex (*β* = .11, *t* = .56, *p* = .58, *β* = .16, *t* = .86, *p* = .39 when controlling for age) interaction effect on AS latency.

### 3.4. Associations between puberty-related fronto-striatal rsFC with antisaccade (AS) performance and latency

#### 3.4.1. Puberty-related effects across males and females on AS

We first examined interactions between puberty and AS performance/latency on rsFC in the fronto-striatal connections that showed significant main effects of puberty after controlling for age^−1^ and sex (i.e., vACC – caudate, dlPFC – NAcc, dlPFC – caudate, vlPFC – NAcc, vlPFC – putamen). Across males and females, dlPFC – caudate rsFC was significantly associated with AS performance as a function of puberty (*β* = .21, *t* = 4.26, *p*_Bonferroni_ = .0001), such that earlier in puberty, stronger rsFC was associated with *worse* performance, and this relationship weakened with further pubertal maturation. In addition, we found that vlPFC – putamen rsFC was significantly associated with AS latency on correct trials as a function of puberty (*β* = -.24, *t* = - 4.30, *p*_Bonferroni_ = .0001), such that earlier in puberty, stronger rsFC was associated with longer latencies (i.e., slower response times) on correct trials, but as pubertal maturation continued, stronger connectivity was associated with shorter latencies (i.e., faster response times). See **Fig. 7** for interaction plots. No other fronto-striatal connection was significantly associated with AS performance or latency on correct trials as a function of puberty (*p*s > .05).

**Figure 7.**
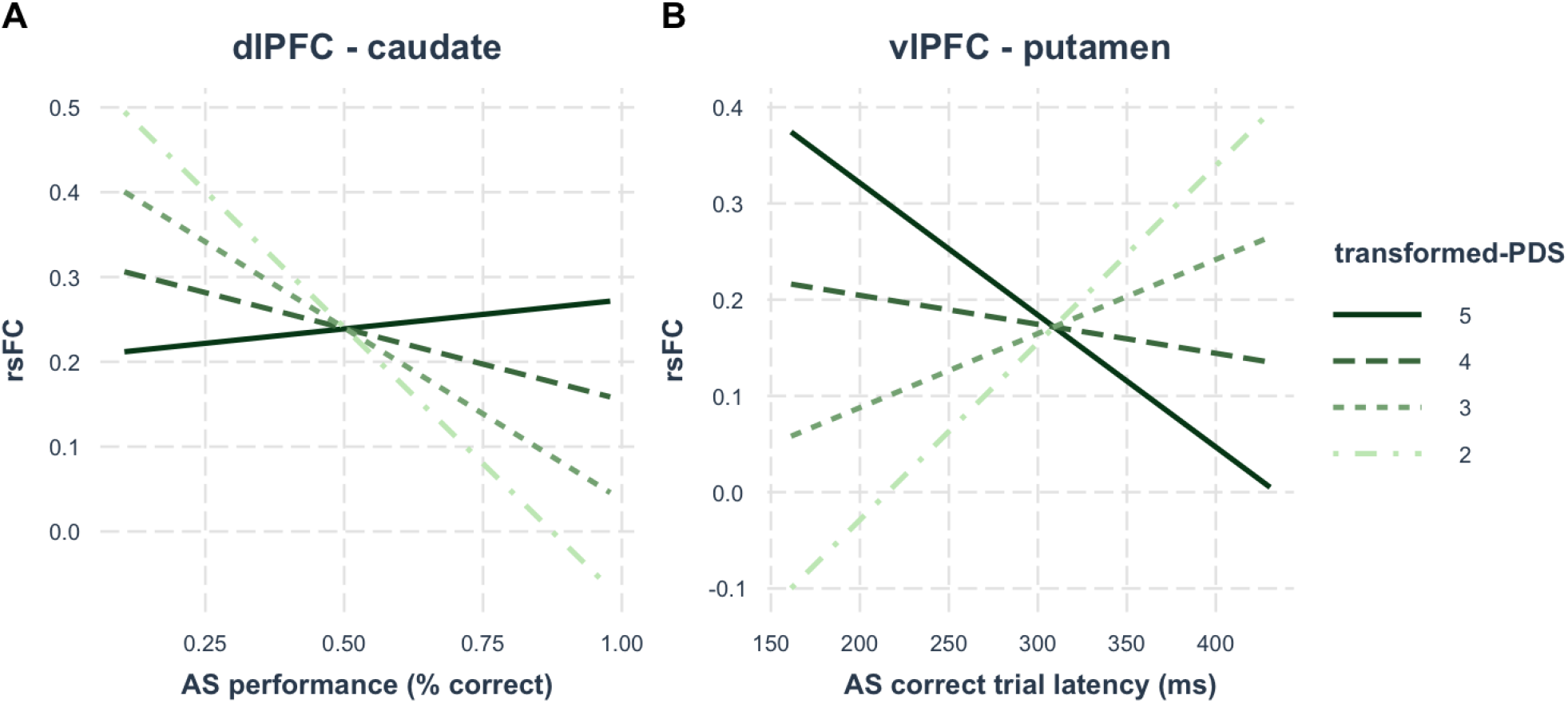
Interaction effects between AS performance/latency and fronto-striatal rsFC across males and females. (**A**) dlPFC – caudate rsFC was associated with AS performance as a function of puberty and (**B**) vlPFC – putamen rsFC was associated with AS correct trial latency as a function of puberty. Abbreivations: dlPFC, dorsolateral prefrontal cortex; vlPFC, ventrolateral prefrontal cortex; rsFC, resting-state functional connectivity; AS, antisaccade; PDS, Petersen Puberty Development Scale.

#### 3.4.2. Sex-specific puberty-related effects on AS

We next tested interactions between puberty and AS performance and latency on rsFC in the fronto-striatal connections that showed significant puberty-by-sex interactions after controlling for age (i.e., anterior vmPFC – caudate, rACC – caudate, vlPFC – caudate), and these analyses were performed separately in males and females. We found that anterior vmPFC – caudate rsFC was significantly associated with AS performance as a function of puberty in both males (*β* = .25, *t* = 3.46, *p*_Bonferroni_ = .004) and females (*β* = -.20, *t* = -2.90, *p*_Bonferroni_ = .025). In males, stronger anterior vmPFC – caudate rsFC was associated with better AS performance, but only in late puberty (approximately following stage 3.4). In contrast, in females, stronger anterior vmPFC – caudate rsFC was associated with better AS performance, but only during early puberty (approximately before stage 2.8). See **Fig. 8** for interaction plots. To formally test whether anterior vmPFC – caudate rsFC was differentially associated with AS performance as a function of sex (between males and females), we ran a post-hoc test with sex modeled as an interaction term. We observed a significant three-way interaction (*β* = .33, *t* = 3.36, *p* = .0009), suggesting that the association between AS performance and anterior vmPFC – caudate rsFC was significantly different between males and females, such that stronger connectivity during early puberty was associated with *better* performance in females but *worse* performance in males. By the end of puberty, however, stronger connectivity was associated with better performance in males but worse performance in females. Finally, in males, we observed significant relationships between AS latency on correct trials and rsFC as a function of puberty for rACC – caudate (*β* = .20, *t* = 2.71, *p*_Bonferroni_ = .043) and vlPFC – caudate (*β* = -.20, *t* = -2.66, *p*_Bonferroni_ = .049). Stronger rACC – caudate rsFC was associated with shorter latencies on correct trials (faster responses) in males during early puberty and this relationship weakened with further pubertal maturation. In contrast, stronger vlPFC – caudate rsFC was associated with longer latencies on correct trials (slower responses) during early puberty, but with shorter latencies on correct trials (faster responses) by late puberty. See **Fig, 9** for interaction plots. No additional fronto-striatal connections were significantly associated with AS performance or latency as a function of puberty (*p*s > .05).

**Figure 8.**
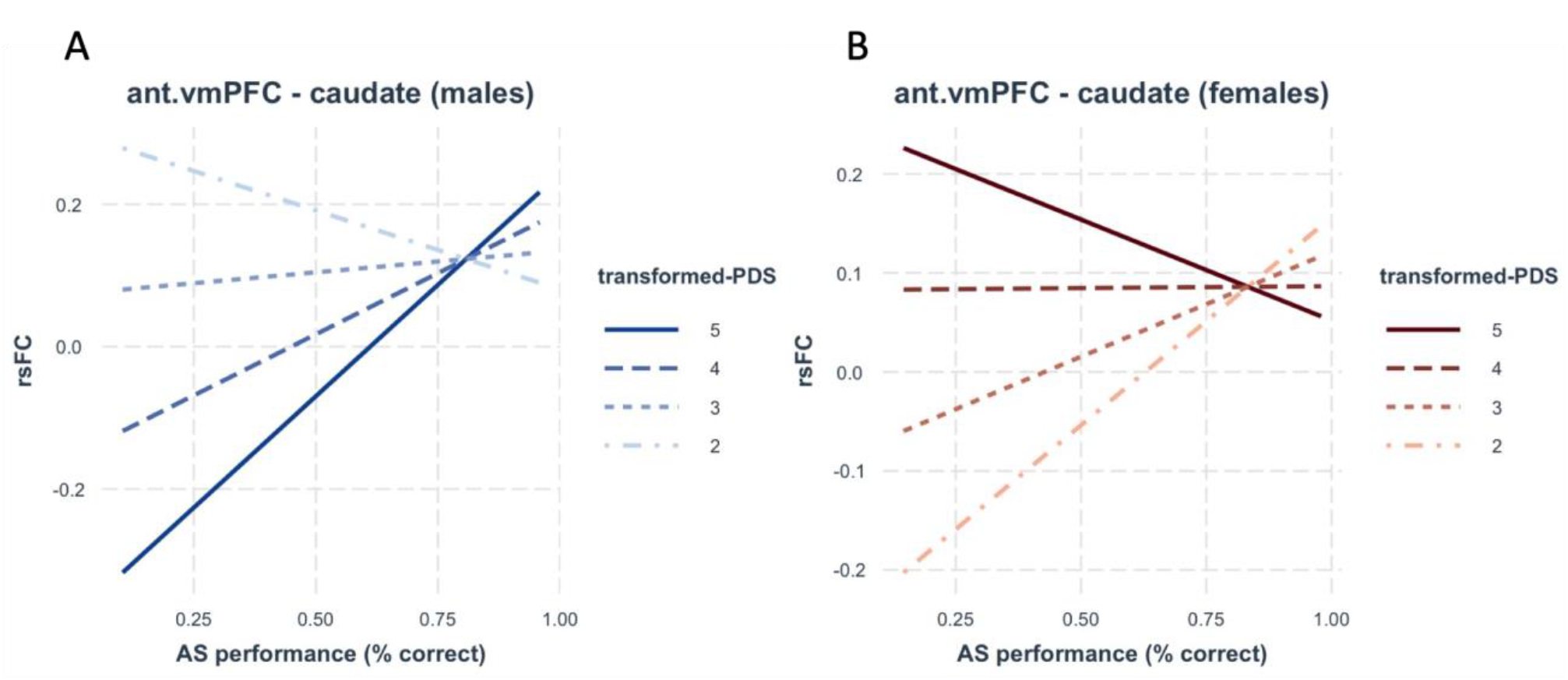
Interaction effects between anterior vmPFC – caudate rsFC and AS performance as a function of puberty in (**A**) males and (**B**) females. Abbreviations: ant.vmPFC, anterior ventromedial prefrontal cortex; rsFC, resting-state functional connectivity; AS, antisaccade; PDS, Petersen Puberty Development Scale.

**Figure 9.**
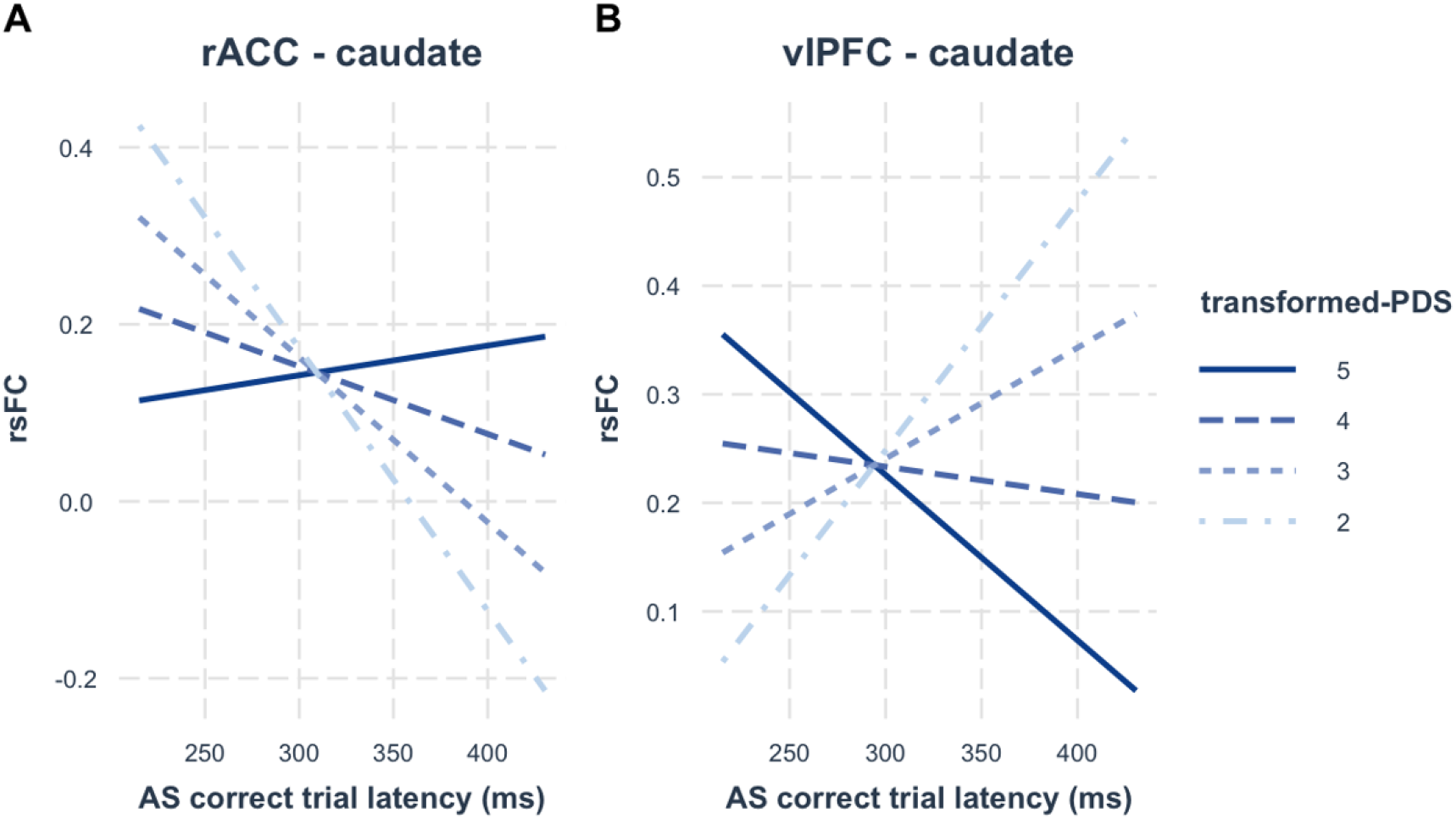
Interaction effects between fronto-striatal rsFC and antisaccade correct trial latency in males as a function of puberty. Interactions were significant for (**A**) rACC – caudate rsFC and for (**B**) vlPFC – caudate rsFC. Abbreviations: rACC, rostral anterior cingulate cortex; vlPFC, ventrolateral prefrontal cortex; rsFC, resting-state functional connectivity; AS, antisaccade; PDS, Petersen Puberty Development Scale.

## Discussion

In this study, we used longitudinal rsfMRI data to investigate the contribution of puberty to fronto-striatal rsFC maturation—over-and-above age-related contributions—and tested associations with inhibitory control using the antisaccade task. Given previous findings indicating puberty-related decreases in rsFC between medial frontal structures and the striatum (van Duijvenvoorde et al., 2019), we hypothesized that puberty would be associated with weaker striatal rsFC with medial frontal regions (i.e., anterior vmPFC, sgACC, rACC, vACC) and stronger striatal rsFC with lateral PFC regions (i.e., dlPFC, vlPFC) after controlling for age-related effects. Our findings partially support this hypothesis: as expected, we did observe puberty-related rsFC decreases between the caudate and anterior vmPFC, rACC, and vACC, however, we also observed puberty-related decreases for dlPFC/vlPFC – caudate and vlPFC – caudate/putamen, which was counter to our initial hypothesis. Further in line with our hypothesis, we observed puberty-related increases in dlPFC – NAcc rsFC. We additionally observed three puberty-by-sex interaction effects on fronto-striatal rsFC in caudate rsFC with the anterior vmPFC, rACC, and vlPFC. Finally, we explored the extent to which the associations between fronto-striatal rsFC and inhibitory responses (performance on the antisaccade task) and efficiency (latency on correct trials) differed as a function of puberty. We found that stronger dlPFC – caudate rsFC during early puberty was associated with worse AS performance in both males and females, and this association became weaker with increased pubertal maturation. Stronger vlPFC – putamen rsFC during early puberty was associated with slower correct responses in both males and females, but by late puberty, stronger vlPFC – putamen rsFC was associated with faster correct responses. We additionally observed that the relationship between anterior vmPFC – caudate rsFC and AS performance differed between males and females as a function of puberty: in males, stronger connectivity was associated with better performance in *late* puberty, but with better performance in females during *early* puberty. To our knowledge, this is the first study to investigate puberty-related fronto-striatal maturation as well as its relationship to the development of inhibitory control.

Interestingly, in comparison to age, we observed a greater number of associations between fronto-striatal connectivity and puberty in adolescence, suggesting that these connections may be specifically influenced by pubertal maturation, whereas others may be independent of pubertal maturation during this period of development. Connections in which we observed puberty-related effects but failed to observe age-related effects (in ages up to 18) included vACC – caudate, dlPFC – NAcc/caudate, and vlPFC – NAcc. Given that our puberty analyses controlled for age-related effects, our findings suggest that puberty may contribute a distinct and dissociable effect on maturation of these fronto-striatal systems during adolescent neurodevelopment. The dlPFC is known to support cognitive processes, including inhibitory control (Angius et al., 2019), working memory (Arnsten and Jin, 2014), action selection (Mars and Grol, 2007), attentional processes (Johnson et al., 2007), and cognitive effort expenditure (Framorando et al., 2021). Previous work, however, has found that dlPFC activation does *not* explain age-related improvements on the antisaccade task (Ordaz et al., 2013), with adolescents and adults exhibiting similar dlPFC recruitment. This may indicate that although local dlPFC functioning may be in place by adolescence, its connectivity to action-related structures (e.g., striatal) may contribute to dlPFC-related cognitive-behavioral improvements, and that this maturation in functional connectivity may be supported by puberty-related processes. Interestingly, age-related effects on dlPFC – NAcc/caudate rsFC, although not observed in our sample prior to age 18, were observed when examining participants through the third decade of life. As such, it may be the case that the puberty effects associated with dlPFC – NAcc/caudate maturation are not yet apparent in samples under 18 but only become evident later in life, possibly bolstered by pubertal maturation; however, *how* puberty might contribute to fronto-striatal maturation (e.g., via hormonal processes) remains unclear. Interestingly, stronger dlPFC – caudate connectivity in earlier pubertal stages, while likely still immature, was associated with *worse* inhibitory control performance, possibly reflecting puberty-related contributions to improvements in inhibitory response control via this circuitry. Indeed, research has implicated dlPFC – caudate functional connectivity in response inhibition (Zhang and Iwaki, 2020) and cognitive control (Yuan et al., 2017). dlPFC – NAcc/caudate rsFC has been implicated in choice impulsivity on a delay-discounting task in a large sample of young adults (n = 429; Wang et al., 2020), which the study authors note they did not observe for the putamen, suggesting specificity of dlPFC – striatal circuitry involvement in more impulsive decisions during earlier stages of maturation but with response inhibition during later stages of development. Our present findings extend our understanding of dlPFC circuitry by implicating puberty as a central process associated with maturation of this system.

As with the dlPFC, the vlPFC also shows age-related maturation through young adulthood (Paulsen et al., 2015), which we similarly observed. Further, our results implicate vlPFC – striatal circuitry in puberty-related neurodevelopment during the adolescent period, as we observed puberty effects with the vlPFC rsFC across all three striatal ROIs beyond age-related effects, possibly indicating that some of the previously reported age-related effects may be capturing puberty-related contributions. Cortical thinning of lateral PFC regions (i.e., dlPFC, vlPFC) during adolescence has been shown to be associated with better cognitive reappraisal—a process of reframing the meaning of affectively charged stimuli to mitigate the emotional impact of said event on the person (Ochsner and Gross, 2008)—but only in females (Vijayakumar et al., 2014), indicating a possible contribution of lateral PFC development to improvements in emotion regulation via cognitive maturation (Phillips et al., 2008; Picó-Pérez et al., 2017; Silvers et al., 2016). We showed a significant puberty-by-sex interaction for vlPFC – caudate rsFC, which was primarily driven by more robust effects observed in females in our sample who exhibited puberty-related rsFC increases following mid-puberty. Further, we observed a significant association between vlPFC – putamen rsFC and inhibitory control efficiency (AS correct trial latency), such that stronger connectivity in late, but not early, puberty was related to faster correct responses. These findings are consistent with previous research that has suggested that the vlPFC may underlie the association between visual cues and action selection (Passingham et al., 2000) and optimal performance (Paulsen et al., 2015). In males, we also observed that stronger vlPFC – caudate rsFC was associated with more efficient inhibitory control (faster correct AS responses), but only in late puberty; interestingly, in early puberty, stronger connectivity was associated with less efficient (slower) correct responses, suggesting that puberty may contribute to vlPFC circuitry maturation supporting efficient inhibitory control. These results suggest that lateral PFC – caudate circuitry related to goal-directed behaviors associated with inhibition and impulsivity (e.g., executing a successful antisaccade response) may improve in a puberty-dependent manner. Taken together our results, show that puberty was predominantly related to changes between prefrontal regions and the caudate, which is linked to supporting cognitive processes (Grahn et al., 2008). The NAcc, which supports reward-processing and motivated behaviors (Murty et al., 2018), was only linked to vl- and dlPFC, which was also evident in its connectivity to the caudate. Whether puberty is *necessary* for the maturation of lateral PFC – striatal systems supporting improvements in cognition, however, remains an open question.

Although we observed a main effect of puberty on medial frontal – striatal regions characterized by rsFC decreases with pubertal maturation, post-hoc tests by sex revealed that in males puberty was associated with weaker rsFC whereas in females further pubertal maturation following mid-puberty (approximately stage 3.5) was associated with *stronger* rsFC. Previous work has shown age-related rsFC increases between medial PFC regions and the caudate (Manza et al., 2015) and separate research has implicated mPFC systems in social closeness and generating affective meaning, which may be particularly relevant during the adolescent period when social relations often become of great significance to adolescents (Krienen et al., 2010; Roy et al., 2012; Somerville et al., 2012). Specifically, in a study of children, adolescents, and young adults, Somerville and colleagues demonstrated using a social-evaluative task that being watched by peers elicited self-conscious emotions (e.g., embarrassment) and physiological arousal (e.g., skin conductance) in adolescents at levels higher than in children or young adults. Further, during task-fMRI, adolescents exhibited stronger mPFC responses during both the anticipation and evaluation phases of the social cognitive paradigm compared to children and young adults, with peak mPFC activation at approximately age 15. Connectivity analyses during task-fMRI further revealed that mPFC – caudate coupling, in particular, during the evaluation phase was significantly stronger in adolescents than in either children or adults, leading the authors to argue that the mPFC might incorporate salient social information (e.g., perceived peer evaluation) with relevance to oneself. Further, they speculated that connectivity with the dorsal striatum (i.e., caudate) might enable integration of relevant social signals to motivational systems, such as those supporting goal-directed behaviors. Although we did not examine social and/or affective neural or behavioral processes in the present study, our post-hoc analyses revealed that caudate rsFC increased during (late) puberty with the anterior vmPFC and rACC only in females. How puberty-related maturation of these mPFC systems relates to affective and social processing in adolescence, however, remains an open question. Given findings indicating that puberty confers greater developmental risk for affective psychopathology during adolescence, particularly in females, atypical maturation of medial PFC – striatal systems may represent one means by which females are at greater risk for developing affective internalizing symptoms (Blair et al., 2013; Greenberg et al., 2013; Ironside et al., 2021), especially in light of evidence linking activation in certain medial frontal structures (e.g., the perigenual ACC and vmPFC) to greater negative self-evaluation in females (Barendse et al., 2020). The specific mechanisms underlying a possible link between the maturation of this circuitry during adolescence to affective and social processes, however, are unclear but likely involve a confluence of developmental changes during puberty, including hormonal, neural, and social factors (Vijayakumar et al., 2018; Ladouceur et al., 2019). As such, future work should investigate puberty-related contributions to such social, affective, and cognitive aspects of adolescent development.

Theories of adolescent development (e.g., The Driven Dual Systems Model) propose an over-engagement of dopaminergic reward systems, which are thought to outpace development of connections from cognitive systems, resulting in a transitionary period biased toward rewarding outcomes and characterized by higher risk-taking and/or sensation-seeking behaviors (Larsen and Luna, 2018; Luciana et al., 2012; Shulman et al., 2016; Spear, 2000). Indeed, evidence from fMRI work supports the mediation of motivated, goal-directed behaviors by way of fronto-striatal networks (Parr et al., 2021). Striatal responses to reward and reward anticipation are more pronounced in adolescents than in either children or adults (Luciana et al., 2012), and evidence suggests that activation in the bilateral caudate and ventral striatum decreases in response to reward anticipation between ages 14 and 19 (Cao et al., 2021). Optimal inhibitory control may be achieved only when there is mature effective coupling with prefrontal systems critical for action selection (e.g., dlPFC) and inhibition (e.g., vlPFC). In contrast, mPFC – striatum functional connectivity—primarily characterized by decreases in connectivity across adolescent development, except for in some cases in females—may be more closely related to social-emotional processes, such as self-consciousness (Somerville et al., 2012), and may mature earlier than lateral PFC systems, as observed in non-human primate studies (Caviness et al., 1995; Orzhekhovskaia, 1977, 1975); however, as we did not directly examine differences in the relative timing of lateral versus medial PFC connections, this hypothesis would require explicit testing.

Our findings are largely consistent with previous puberty-related functional connectivity evidence previously reported. For example, in a well-powered study sample studied longitudinally, van Duijvenvoorde and colleagues observed puberty-related decreases in rsFC between striatal regions and frontal medial and ACC subregions (2019). Previous work using structural MRI evidence has observed puberty-related volumetric *decreases* in the NAcc (Goddings et al., 2014) and evidence from animal research indicates that androgens and estrogens influence dopamine release in the NAcc (Thompson and Moss, 1994), hinting at one possible mechanism by which puberty exerts its neurodevelopmental effects on striatal maturation. Although specific mechanisms remain unknown, investigators have posited that a confluence of exposure to hormones (e.g., testosterone, estradiol) and psychosocial stress and stimulation (e.g., new romantic interests, academic or work-related demands) may act in conjunction during this critical period of cortico-subcortical (e.g., fronto-striatal) neurodevelopment (Byrne et al., 2017; Larsen and Luna, 2018; Sinclair et al., 2014). Previous work has implicated puberty—and sex hormones, in particular—as a core mechanism contributing to adolescent cortical maturation (Delevich et al., 2021; Drzewiecki et al., 2016). However, given the lack of studies investigating neuroendocrinological development in typically developing human adolescents, it remains difficult to identify the exact mechanisms and pathways underlying puberty-related maturation of neurobiological systems associated with cognitive and affective developmental processes. Several lines of evidence implicate dopaminergic systems as one key link between puberty—and gonadal sex hormones, in particular—and neurodevelopment supporting cognitive maturation (Hernandez et al., 1994; Kuhn et al., 2009; Ladouceur et al., 2018). Findings from experiments carried out in animal models suggest that puberty and dopaminergic neurophysiological maturation not only overlap in their developmental timelines, but center puberty as a potentially necessary factor for maturation of dopaminergic circuitry and goal-directed behaviors (Bell et al., 2013). Animal models have shown that dopamine can inhibit pubertal hormone synthesis (i.e., luteinizing hormone), and increased dopamine receptor binding correlates with testosterone levels, particularly in females (Andersen et al., 2002), suggesting an intricate interplay between these two systems to support the transition from adolescence to adulthood (Vidal et al., 2004). Further, animal research indicates that sex steroids contribute to the reorganization and specialization of neural circuitry via the effects of hormones on synapses (Fernandez–Galaz et al., 1997; Matsumoto, 1991; Parducz et al., 2006), thereby specializing neural connections—via synaptic pruning and synaptogenesis in various structures, for example—to support adult-like behaviors (Schulz and Sisk, 2016).

Our current findings advance our understanding of the development of fronto-striatal systems, by indicating that pubertal maturation, especially around mid-puberty when gonadal hormones reach their highest concentrations (Balzer et al., 2019), may underlie some of the developmental neurobiological and behavioral effects observed in the maturation of this system. Our results propose that integration between executive lateral prefrontal regions involved in action-selection and inhibitory control (i.e., dlPFC, vlPFC) and striatal regions supporting cognitive (caudate) and motivational (NAcc) processes may undergo specific refinement guided by pubertal maturation that may define their stability in adulthood. Our work suggests that an inflection point, around mid-puberty, may be crucial in initiating the “beginning of the end” of the adolescent critical period, when cognitive processes more closely resemble adult-like behaviors.

### Study limitations and strengths

Our findings should be considered in light of several limitations. First, our puberty measure was self-reported. Evidence suggests that although self-report measures and clinical physical assessment correspond, there are some caveats in using only the former: specifically, whereas boys tend to report being more pubertally advanced compared to clinical assessments, girls tend to underreport their pubertal stage (Rasmussen et al., 2014). Critically, we also only examined associations in individuals in whom pubertal maturation was already underway (transformed-PDS ≥ 2) and our sample was negatively skewed, such that pubertally mature adolescents were overrepresented, which may limit our ability to detect and interpret effects associated with earlier periods of adolescence and puberty. As such, we are cautious in characterizing these associations between fronto-striatal rsFC and puberty as reflecting a continuous relationship throughout the entire pubertal maturation period. Rather, we believe our findings mainly speak to pubertal maturation from stages when pubertal maturation is underway (i.e., stage ≥ 2), and, more so, to the middle-to-end of pubertal maturation where our sample is most strongly powered to detect changes. Further, some maturational effects captured in older participants (i.e., ages 10 and older) are likely more prominent in female participants as compared to male participants, given earlier pubertal maturation in females (Patton and Viner, 2007). We also did not administer the PDS to participants over the age of 18, thereby precluding characterization of the end of pubertal development for some in our sample who had not reached pubertal maturation (i.e., stage 5) by this age, particularly for males. Ideally, studies examining pubertal maturation should consider recruiting participants at earlier stages before puberty is underway and continuing to administer pubertal assessments until adult-like maturation (i.e., stage 5) is reached. Our study leveraged a large, longitudinal study design and used naturally occurring variability in age and puberty to disentangle their effects. This approach requires large samples to achieve sufficient power to fully characterize each. An alternative approach is to recruit a more age-homogenous sample, or to specifically recruit based on age and pubertal status in order to maximize variability between pubertal maturation and age (which may also differ as a function of sex, given sex-related differences in pubertal timing). This approach has the advantage of removing the need to regress out age effects and allows better powered analyses for age versus puberty in smaller samples. Notably, this approach has the disadvantage of having to limit the age range (e.g., only recruiting between ages 10-12 or 16-18, etc.), or oversampling participants who are pubertally mature or immature for their age. Future work that leverages extremely large, longitudinal, multimodal datasets—such as those collected in the Adolescent Brain Cognitive Development (ABCD) study—may provide the best opportunity to disentangle the highly correlated age and puberty terms, by providing substantial statistical power with non-biased sampling strategies. Further, we were underpowered to test in our models *change* in pubertal maturation intra-individually (i.e., visit-to-visit in one person), which may further reveal important associations with brain maturation. Experimenters investigating puberty might consider widening the age range of participants during initial recruitment and continue to assess pubertal maturation at every visit until participants reach adult-like maturation (i.e., Tanner Stage 5). Multiple puberty datapoints on each participant would enable modeling of pubertal *tempo*, or the rate at which one advances pubertally, which may be more closely associated with certain neural systems and behaviors. Finally, we were unable to include random slopes in our models due to most participants having two or fewer longitudinal rsfMRI visits and pubertal data. Additional repeated measures of this sort would allow for a more comprehensive characterization of individual differences in trajectories of neurocognitive development.

In addition, our rsfMRI data from the CogLong sample were extracted from blocked design task-fMRI during fixation periods, not from a continuously collected rsfMRI scan, which is an important difference to consider; however, previous evidence has noted such methods to be effective for conducting rsFC analyses (Fair et al., 2007). Although our CogLong rsfMRI data was approximately 6min48s in total, it was collected over an 18-minute period: previous work has indicated that rsFC correlation estimates stabilize in scan times as short as 5min, produce moderate to high test-retest reliability, and are more robust when participants are presented with a fixation and keep their eyes open during scan acquisitions (Dijk et al., 2010), both of which were the case in our sample. Despite differences in acquisition time, our rsFC analyses showed similar age-related effects across ROI pairs to what has been reported in the literature with similar (van Duijvenvoorde et al., 2019; Fareri et al., 2015) and longer (Parr et al., 2021) acquisition times. As such, the fact that our analyses recapitulate prior age-related findings indicates that we should be well positioned to probe pubertal effects (above and beyond age). Nevertheless, future studies should consider lengthening rsfMRI scans, which seems to improve intrasession ICCs until plateauing around 13 minutes (Birn et al., 2013). Longer scan times (i.e., 15-25 minutes), however, are likely required to differentiate an individual’s rsfMRI “fingerprint” from a control group (Anderson et al., 2011). Finally, most developmental cognitive neuroscience studies to date, including ours, have not considered gender identity as it pertains to adolescent development—in contrast to sex assigned at birth—thereby precluding characterizations of cognitive and affective neural and behavioral development more comprehensively across various populations of adolescents. Given the acute risk of affective symptomatology (i.e., depression, anxiety, suicidality) associated with puberty in transgender, non-binary, and/or gender non-conforming children and adolescents—especially when prevented from receiving gender-affirming medical and psychological care (Tordoff et al., 2022; Green et al., 2022; Turban et al., 2022)—we recommend future studies include measures, such as self-report questionnaires, to further advance our understanding of adolescent development.

Despite these limitations, our study has several considerable strengths. First, we leveraged two large, longitudinal, multimodal datasets to test linear and non-linear pubertal contributions to fronto-striatal functional connectivity. We focused our analyses on pubertal maturation by only including participants with PDS ≥ 2 to ensure that the maturational processes of interest were already underway. Additionally, our large extended (full) sample (ages 8 – 34) with several longitudinal timepoints allowed us to track age-related effects through adolescence and into adulthood thereby situating our findings in the context of adult fronto-striatal circuitry more broadly. This is crucial given that we observed several significant age-related associations with fronto-striatal rsFC in this extended sample, which we did not observe when only examining effects in ages up to 18. Finally, by examining both medial and lateral PFC subregional connections with the NAcc, caudate, and putamen, we comprehensively characterize for the first time, to our knowledge, the contributions of puberty to fronto-striatal circuity development, which, in turn, supports cognitive processes such as inhibitory control during adolescence.

### Conclusions & future directions

Our findings add to a nascent yet growing body of scientific literature investigating the effects pubertal maturation might have on the developing adolescent brain. Notably, our results highlight specific fronto-striatal connections that may be influenced by puberty-related processes. This may suggest that some fronto-striatal maturational effects are more specific to pubertal processes than are others. These results also suggest that maturation related to puberty may add specific specialization supporting later maturation into adulthood. Finally, given the sex specificity of some of our findings, we believe this work may inform differential sex-related risks of psychopathology that emerge during this time, especially as they relate to fronto-striatal circuitry, as well as transdiagnostic phenotypes related to inhibitory control and/or impulsivity.

Puberty-related connections identified here may represent a potential therapeutic target, given the relationship between adolescent inhibitory control and substance use vulnerability (Quach et al., 2020), and relationships between fronto-striatal connectivity and substance dependency (i.e., dlPFC – caudate; Qian et al., 2020; Ma et al., 2018). Preliminary findings using transcranial magnetic stimulation (TMS) to target this circuitry indicate promising outcomes for alleviating craving-related symptoms across populations and substances (Hanlon et al., 2015). Therefore, our work may inform timing and targets for preventative strategies during one of the most formative periods of human development.

## Supplemental Materials

### Supplemental Results

#### Associations between age and pubertal maturation

Linear regressions showed that age and pubertal maturation were highly correlated in both males (*r* = .80, *p* < .001, 95%CI[.72, .85]) and females (*r* = .77, *p* < .001, 95%CI[.68, .84]).

#### Sensitivity analyses controlling for motion

To ensure that our primary analyses were not biased by head motion, we additionally controlled for the percent of TRs censored for each subject for all significant associations between puberty and fronto-striatal rsFC. All significant associations remained significant after including this additional covariate (see **Table S1**), which we would expect given our exclusion of functional volumes with a frame-wide displacement (FD) > 0.3 mm along with a scan being entirely excluded from analyses if over 40% of volumes were censored.

#### Seed-based approach

No cluster reached statistical significance (*p* < .01) at an alpha = .05 when we tested for associations with puberty and rsFC between any of the striatal seed ROIs and clusters across frontal regions.

#### Age effects on fronto-striatal rsFC

##### Main effects of age (up to 18-years-old)

Across males and females, we observed significant age-related increases in several fronto-striatal connections, including the anterior vmPFC – NAcc (*F* = 7.10, *p*_Bonferroni_ = .016), anterior vmPFC – caudate (*F* = 6.58, *p*_Bonferroni_ = .026), and rACC – caudate (*F* = 6.65, *p*_Bonferroni_ = .024). We additionally observed age-related decreases in two connections: vlPFC – caudate (*F* = 7.14, *p*_Bonferroni_ = .015) and vlPFC – putamen (*F* = 12.9, *p*_Bonferroni_ = .0001). No other fronto-striatal connections examined were significantly associated with age (*p*s > .05).

##### Age-by-sex interaction effects (up to 18-years-old)

We observed three significant age-by-sex interactions: anterior vmPFC – caudate (*F* = 7.87, *p*_Bonferroni_ = .007), rACC – caudate (*F* = 24.89, *p*_Bonferroni_ < .0001), and vACC – caudate (*F* = 11.344, *p* = .0002). Post-hoc tests revealed significant age-related increases in males across all three connections: anterior vmPFC – caudate (*F* = 24.89, *p*_Bonferroni_ < .0001), rACC – caudate (*F* = 24.89, *p*_Bonferroni_ < .0001), and vACC – caudate (*F* = 24.89, *p*_Bonferroni_ = .004); however, we found no significant effect of age in females across these connections (*p*s > .05).

##### Main effects of age (up to 35-years-old)

Across males and females, we observed significant age-related effects for the following fronto-striatal connections: anterior vmPFC – caudate (*F* = 31.48, *p*_Bonferroni_ < .0001), anterior vmPFC – putamen (*F* = 6.21, *p*_Bonferroni_ = .036), sgACC – NAcc (*F* = 16.77, *p*_Bonferroni_ < .0001), rACC – caudate (*F* = 16.18, *p*_Bonferroni_ < .0001), dlPFC – NAcc (*F* = 16.03, *p*_Bonferroni_ < .0001), dlPFC – putamen (*F* = 17.54, *p*_Bonferroni_ < .0001), and vlPFC – putamen (*F* = 7.40, *p*_Bonferroni_ = .0108). All connections, except for sgACC – NAcc and vlPFC – putamen, exhibited age-related increases until approximately age 20 before stabilizing. In contrast, vlPFC – putamen rsFC decreased until age 20 before stabilizing, while sgACC – NAcc rsFC decreased linearly across age. No other connections were significantly associated with age (*p*s > .05).

##### Age-by-sex interaction effects (up to 35-years-old)

In addition, we observed significant age-by-sex interactions on fronto-striatal rsFC for the following connections: anterior vmPFC – NAcc (*F* = 21.14, *p*_Bonferroni_ < .0001), anterior vmPFC – caudate (*F* = 22.24, *p* < .0001), anterior vmPFC – putamen (*F* = 7.91, *p* = .007), rACC – NAcc (*F* = 18.01, *p*_Bonferroni_ < .0001), rACC – caudate (*F* = 11.04, *p*_Bonferroni_ = .0003), and vACC – NAcc (*F* = 6.88, *p*_Bonferroni_ = .019). Post-hoc tests performed separately in males and females revealed several sex-specific effects. In males, rsFC significantly increased with age in the following connections: anterior vmPFC – NAcc (*F* = 10.47, *p*_Bonferroni_ = .0002), anterior vmPFC – caudate (*F* = 48.28, *p*_Bonferroni_ < .0001), anterior vmPFC – putamen (*F* = 10.52, *p*_Bonferroni_ = .0002), rACC – NAcc (*F* = 6.63, *p*_Bonferroni_ = .008), and rACC – caudate (*F* = 25.23, *p*_Bonferroni_ < .0001). In females, rsFC significantly increased until approximately age 20 before decreasing, exhibiting quadratic (inverse-U) shaped trajectories in the following three connections: anterior vmPFC – NAcc (*F* = 16.59, *p*_Bonferroni_ < .0001), anterior vmPFC – caudate (*F* = 6.07, *p*_Bonferroni_ = .012), and rACC – NAcc (*F* = 13.18, *p*_Bonferroni_ < .0001). No other connections were significantly associated with age (*p*s > .05).

#### Developmental effects on antisaccade (AS) performance and latency

##### Associations with puberty

AS performance (i.e., proportion of correct trials) significantly improved with pubertal maturation after covarying for sex (*β* = .33, *t* = 3.94, *p* < .001), but not after also controlling for age (*β* = .02, *t* = .17, *p* = .861), consistent with previous findings (Ordaz et al., 2017). We also failed to observe a significant puberty-by-sex interaction effect (*β* = .20, *t* = 1.25, *p* = .213) on AS performance.

AS latency (on correct trials) significantly decreased (i.e., grew shorter in time) as pubertal maturation advanced (*β* = -.24, *t* = -2.51, *p* = .013) after controlling for sex, but not after controlling for age as well (*β* = .13, *t* = .90, *p* = .371). We also failed to observe a significant puberty-by-sex interaction effect on AS latency (*β* = .16, *t* =.86, *p* = .394).

##### Associations with age (up to 18)

AS performance significantly improved with age with sex modeled as a covariate (*β* = .50, *t* = 7.78, *p* < .001). We failed to observe a significant age-by-sex interaction effect on AS performance (*β* = .12, *t* = .91, *p* = .366).

AS latency significantly decreased with age with sex modeled as a covariate (*β* = -.58, *t* = -8.68, *p* < .001). We also failed to observe a significant age-by-sex interaction effect on AS latency (*β* = .04, *t* = .31, *p* = .759).

##### Associations with age (up to 35)

AS performance significantly improved with age with sex modeled as a covariate (*β* = .32, *t* = 7.19, *p* < .001), but we again did not observe a significant age-by-sex interaction effect (*β* = .01, *t* = .16, *p* = .871).

AS latency significantly decreased with age with sex modeled as a covariate (*β* = -.48, *t* = -11.19, *p* < .001). We again failed to observe a significant age-by-sex interaction effect (*β* = .04, *t* = .51, *p* = .613).

#### Variance of outcome measures (fronto-striatal rsFC, antisaccade performance/latency)

Plots illustrating the variance of our outcome measures of interest are presented in **Fig. S9-10**.

#### Antisaccade performance and latency across puberty and age by trial type

Plots depicting the latency and proportion of antisaccade trials by trial type (i.e., dropped, error, error-corrected, correct) are now presented across age and puberty in **Fig. S11-12**.

## Supplemental Figures

**Figure S1.**
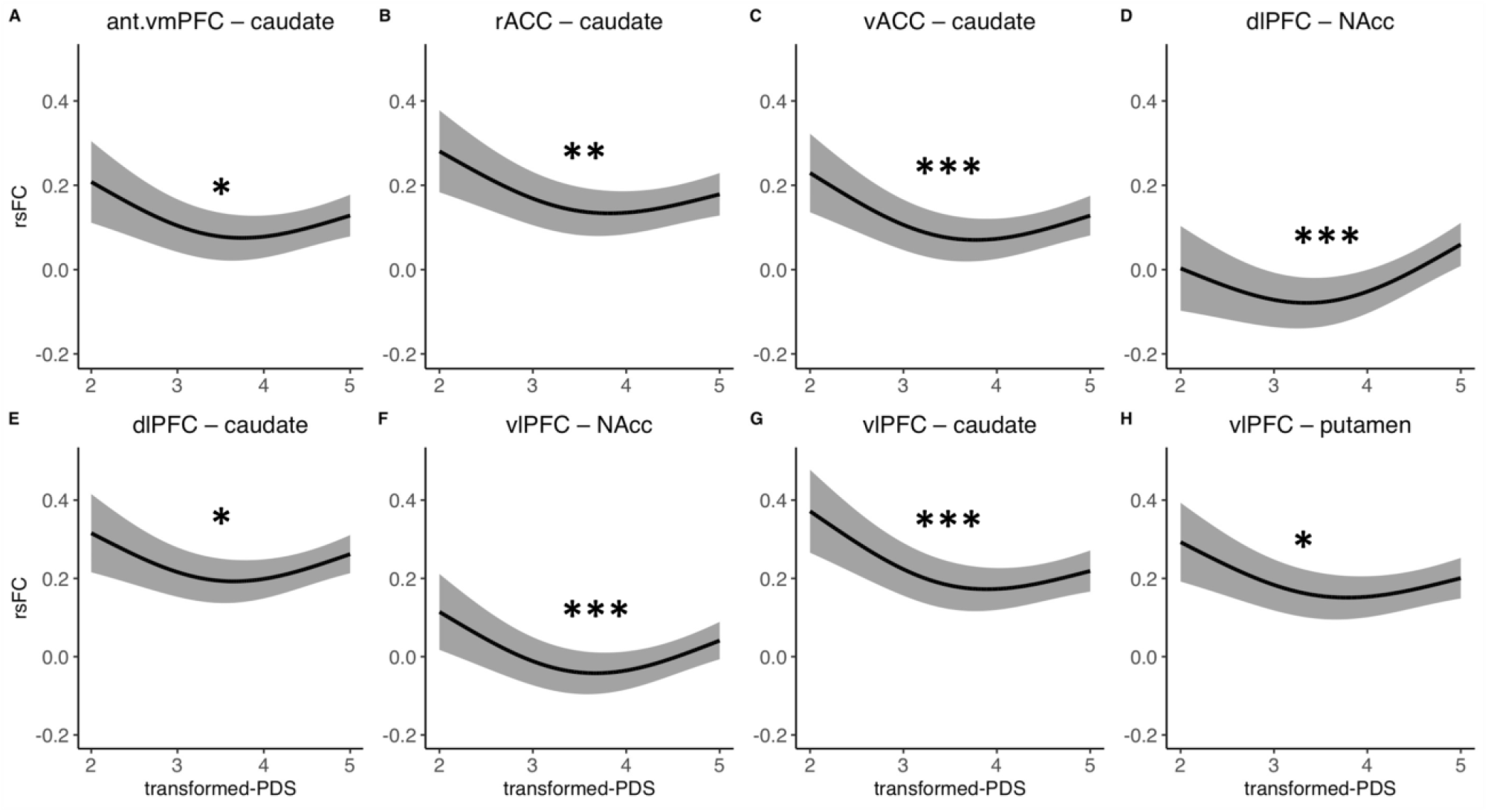
Main effects of puberty on fronto-striatal resting-state functional connectivity (rsFC) beyond age and sex effects. Abbreviations: NAcc, nucleus accumbens; vmPFC, ventromedial prefrontal cortex; sgACC, subgenual cingulate; vACC, ventral anterior cingulate cortex; rACC, rostral anterior cingulate cortex; dlPFC, dorsolateral prefrontal cortex; vlPFC, ventrolateral prefrontal cortex; PDS, Petersen Pubertal Development Scale. * *p*_Bonferroni_ < .05, ** *p*_Bonferroni_ < .01, *** *p*_Bonferroni_ < .001.

**Figure S2.**
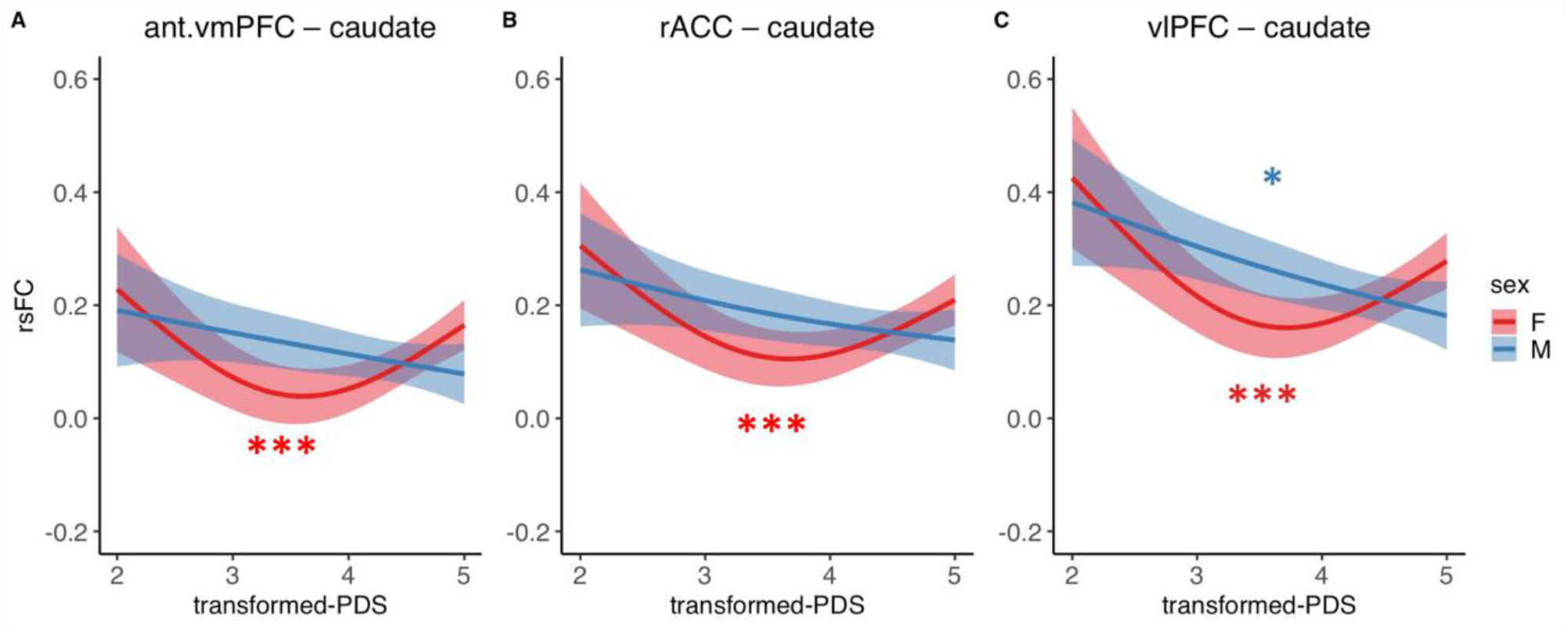
Puberty-by-sex interaction effects beyond age on fronto-striatal rsFC. Asterisks denoting significance refer to post-hoc within sex puberty effects. Abbreviations: ant.vmPFC, anterior ventromedial prefrontal cortex; rACC, rostral anterior cingulate cortex; vlPFC, ventrolateral prefrontal cortex; PDS, Petersen Pubertal Developmental Scale; rsFC, resting-state functional connectivity. * *p*_Bonferroni_ < .05, ** *p*_Bonferroni_ < .01, *** *p*_Bonferroni_ < .001.

**Figure S3.**
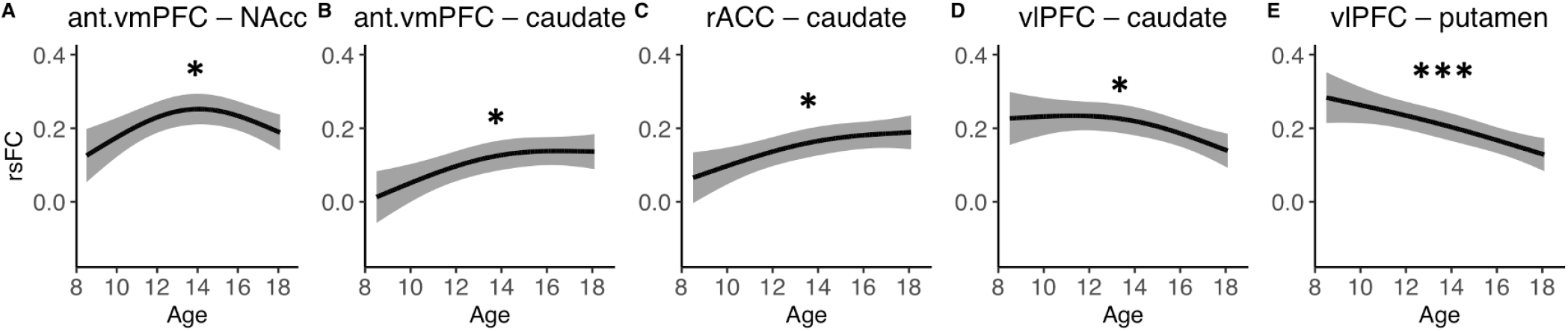
Main effects of age (up to 18-years-old) on fronto-striatal resting-state functional connectivity (rsFC) with sex modeled as a covariate. Abbreviations: ant.vmPFC, anterior ventromedial prefrontal cortex; rACC, rostral anterior cingulate cortex; vlPFC, ventrolateral prefrontal cortex; rsFC, resting-state functional connectivity. * *p*_Bonferroni_ < .05, ** *p*_Bonferroni_ < .01, *** *p*_Bonferroni_ < .001.

**Figure S4.**
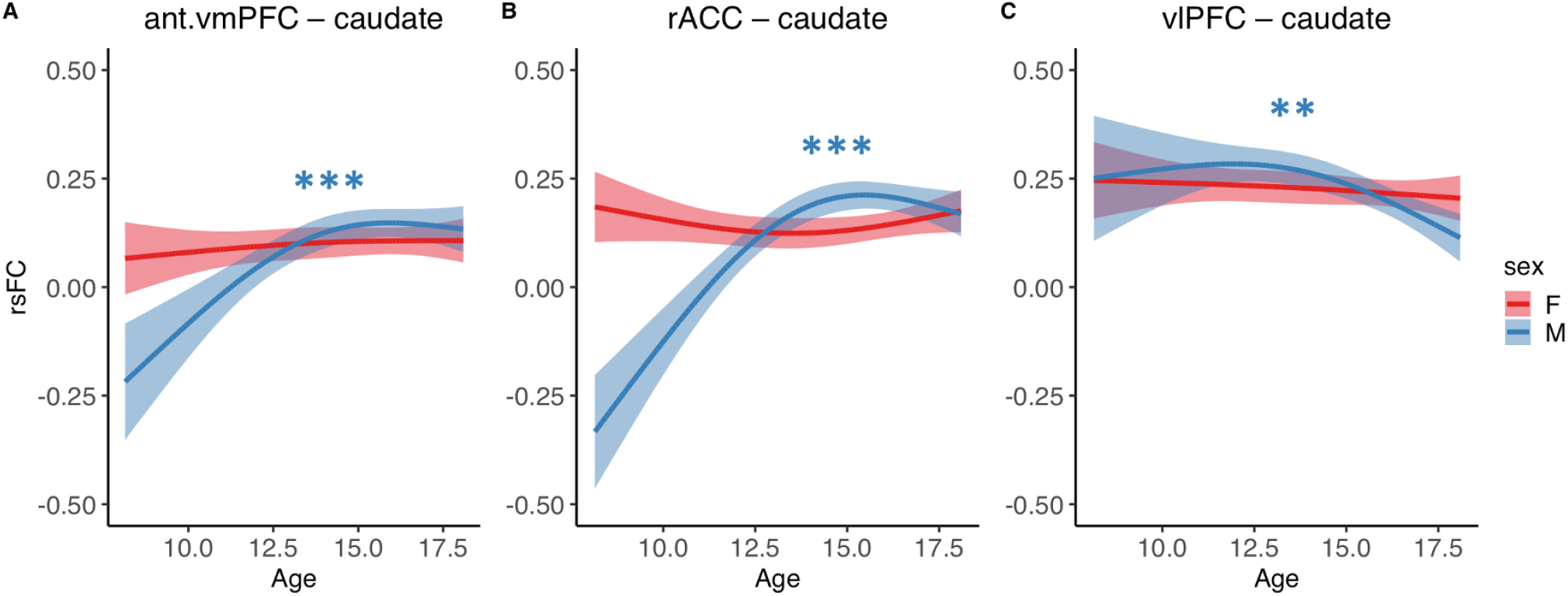
Age-by-sex interaction effects (up to 18-years-old) on fronto-striatal resting-state functional connectivity (rsFC). Asterisks denoting significance refer to post-hoc within sex puberty effects. Abbreviations: ant.vmPFC, anterior ventromedial prefrontal cortex; rACC, rostral anterior cingulate cortex; vlPFC, ventrolateral prefrontal cortex; rsFC, resting-state functional connectivity. * *p*_Bonferroni_ < .05, ** *p*_Bonferroni_ < .01, *** *p*_Bonferroni_ < .001.

**Figure S5.**
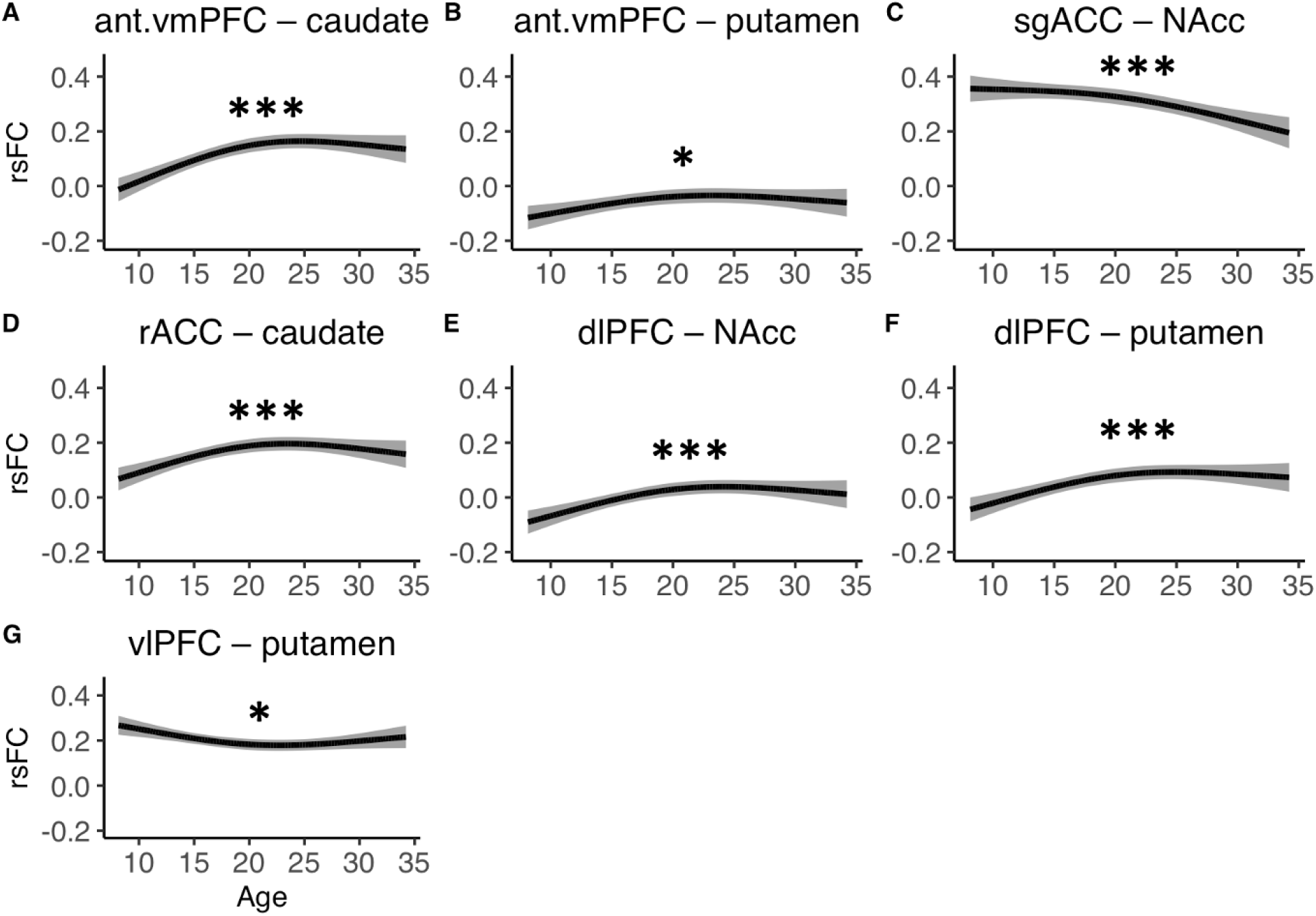
Main effects of age (up to 35-years-old) on fronto-striatal resting-state functional connectivity (rsFC) with sex modeled as a covariate. Abbreviations: ant.vmPFC, anterior ventromedial prefrontal cortex; sgACC, subgenual anterior cingulate cortex; rACC, rostral anterior cingulate cortex; dlPFC, dorsolateral prefrontal cortex; vlPFC, ventrolateral prefrontal cortex; NAcc, nucleus accumbens; rsFC, resting-state functional connectivity. * *p*_Bonferroni_ < .05, ** *p*_Bonferroni_ < .01, *** *p*_Bonferroni_ < .001.

**Figure S6.**
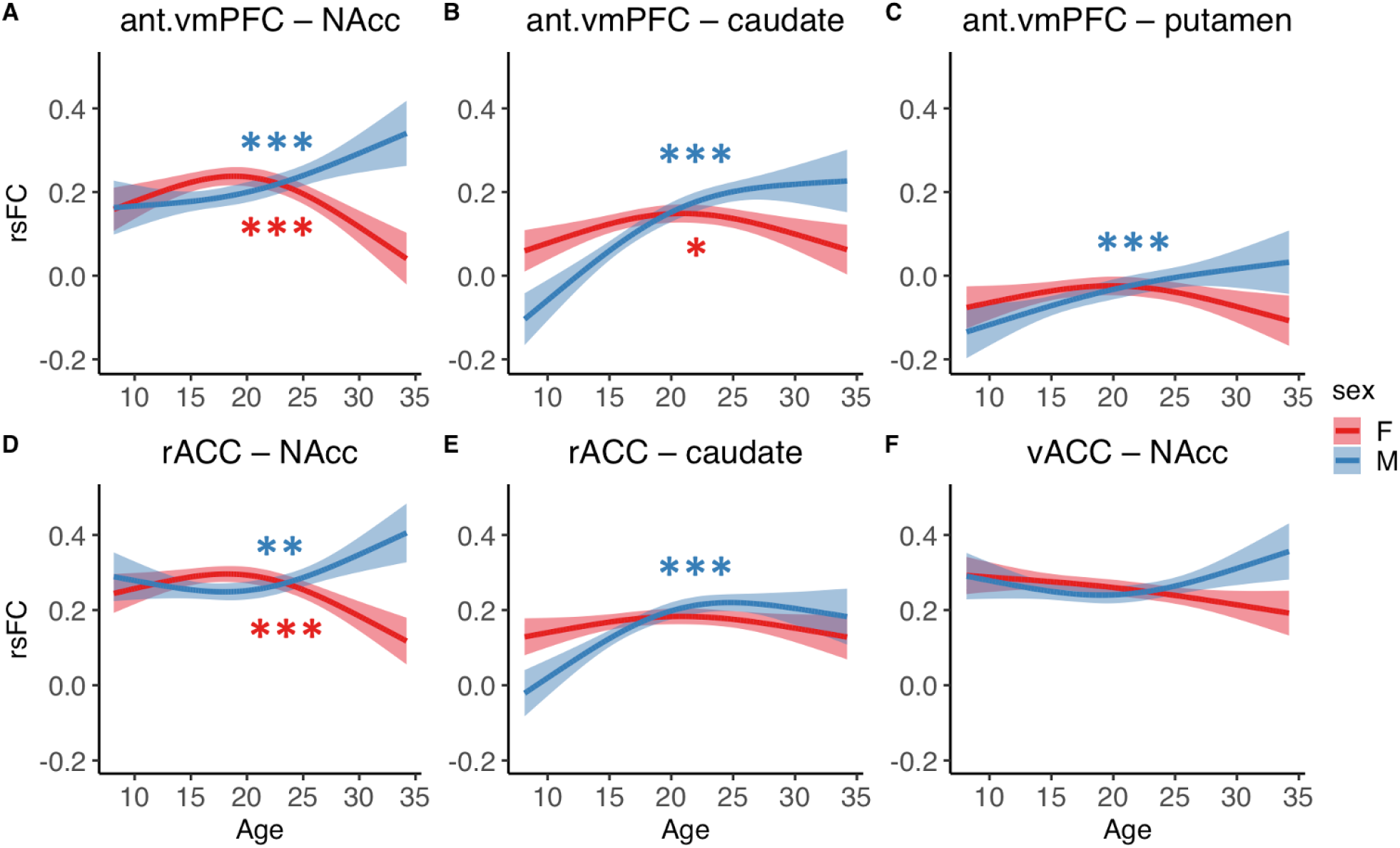
Age-by-sex interaction effects (up to 35-years-old) on fronto-striatal resting-state functional connectivity (rsFC). Asterisks denoting significance refer to post-hoc within sex puberty effects. Abbreviations: ant.vmPFC, anterior ventromedial prefrontal cortex; rACC, rostral anterior cingulate cortex; vACC, ventral anterior cingulate cortex; NAcc, nucleus accumbens; rsFC, resting-state functional connectivity. * *p*_Bonferroni_ < .05, ** *p*_Bonferroni_ < .01, *** *p*_Bonferroni_ < .001.

**Figure S7.**
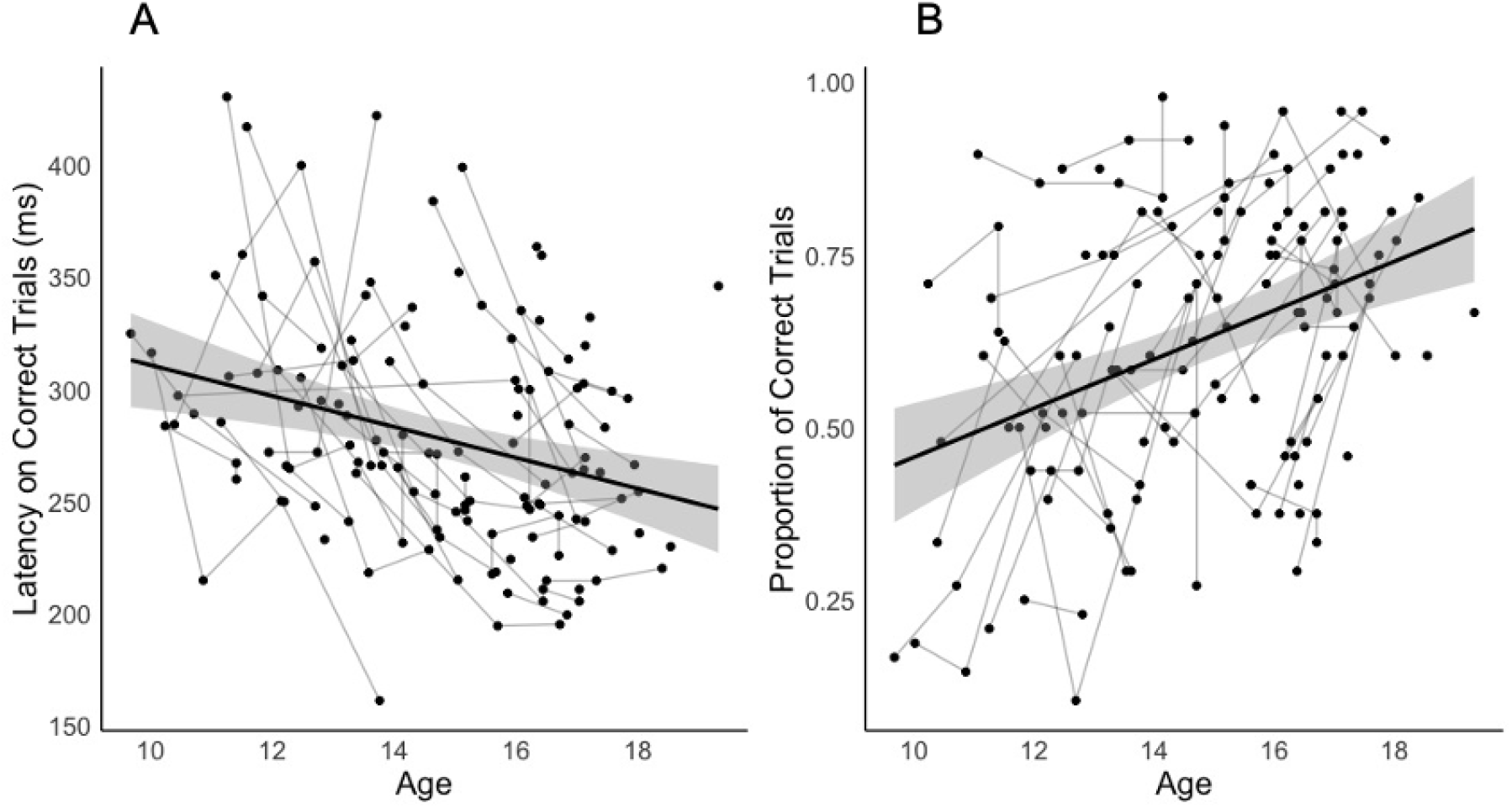
Antisaccade task performance on correct trials by age (up to 18-years-old) separated by (**A**) latency on correct trials and (**B**) proportion of correct trials. Each point indicates a visit and lines indicate multiple visits by the same participant.

**Figure S8.**
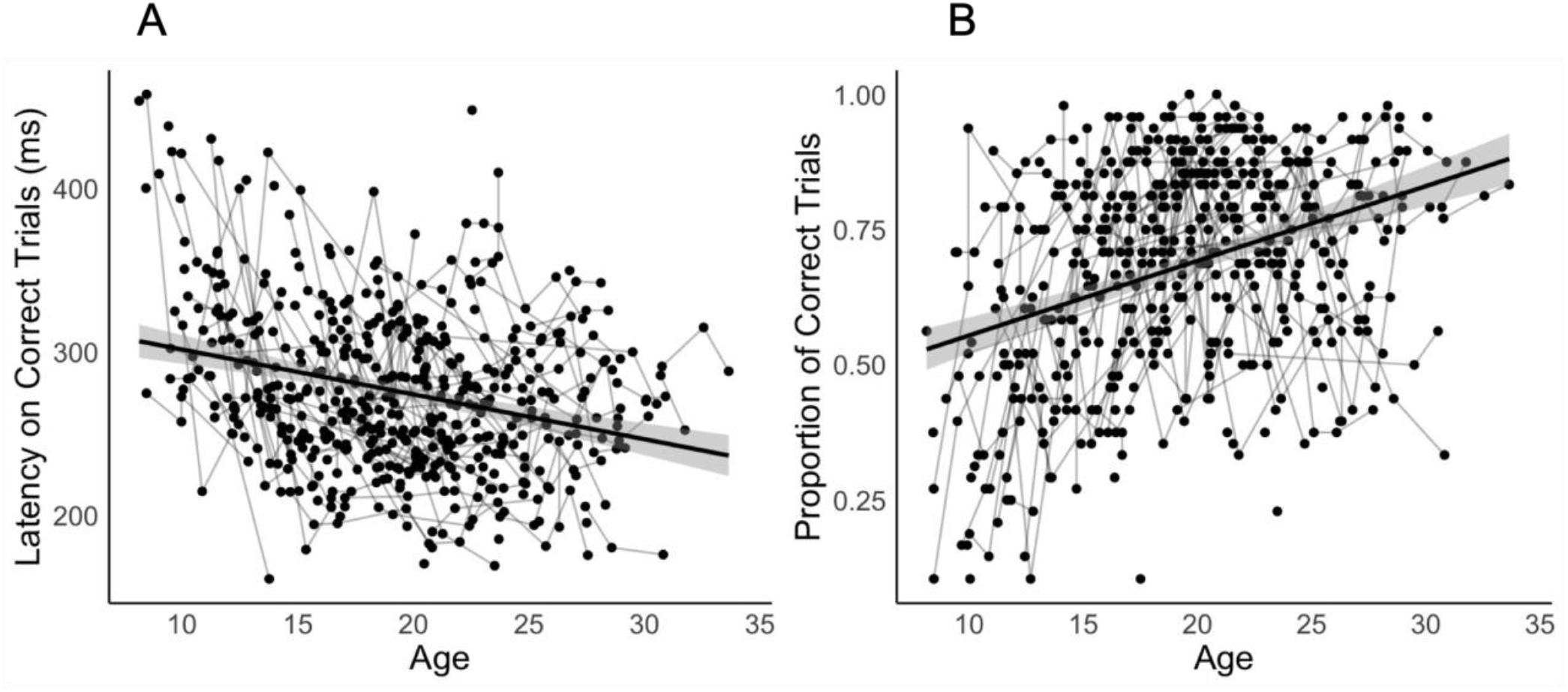
Antisaccade task performance on correct trials in full (up to 35-years-old) age sample by (**A**) latency on correct trials and (**B**) proportion of correct trials. Each point indicates a visit and lines indicate multiple visits by the same participant.

**Figure S9.**
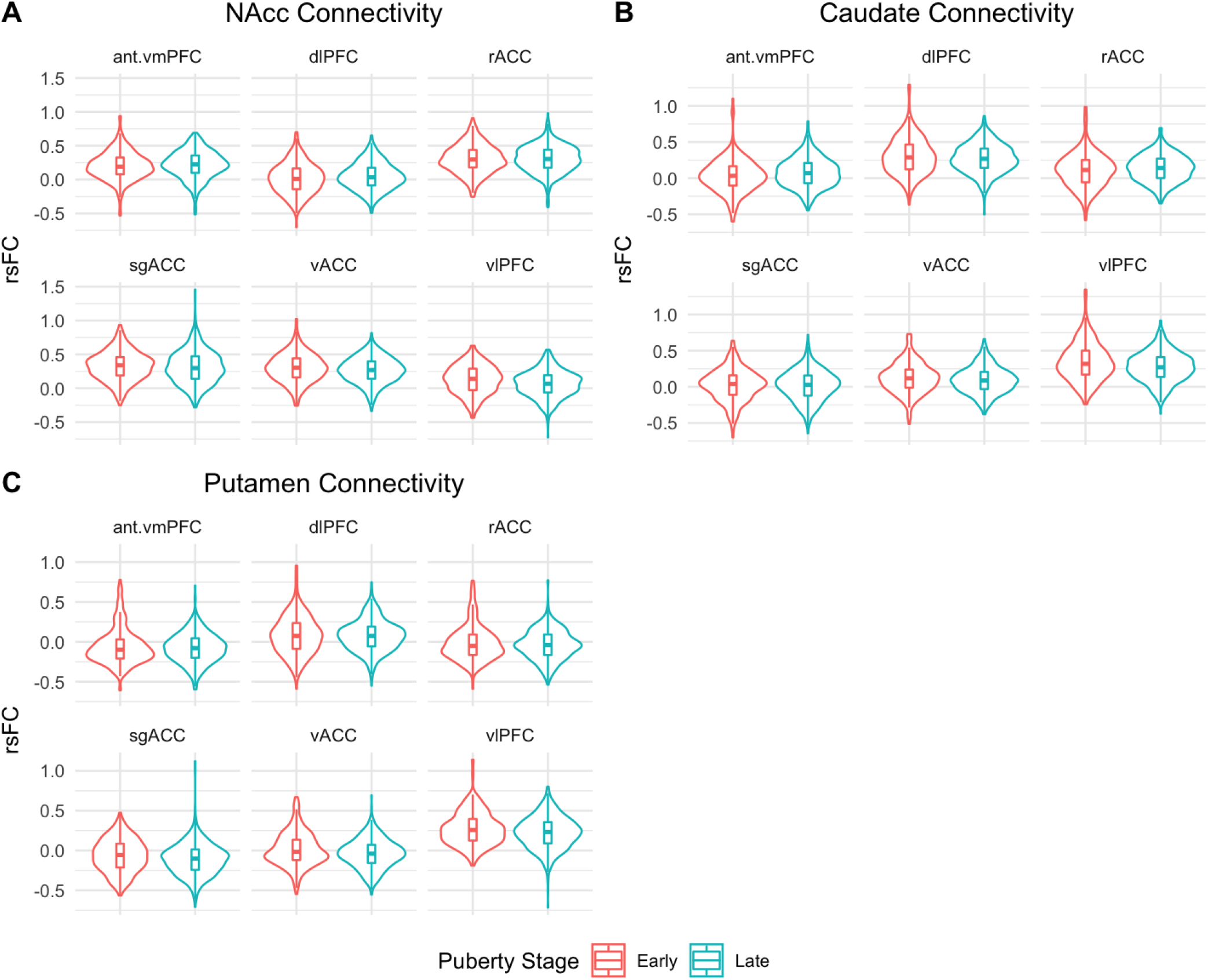
Distribution of fronto-striatal rsFC organized by striatal ROIs: (**A**) NAcc connectivity, (**B**) caudate connectivity, (**C**) putamen connectivity. ‘Early’ puberty stage is defined in the above plots as transformed-PDS < 4 and ‘Late’ puberty stage as transformed-PDS ≥ 4.

**Figure S10.**
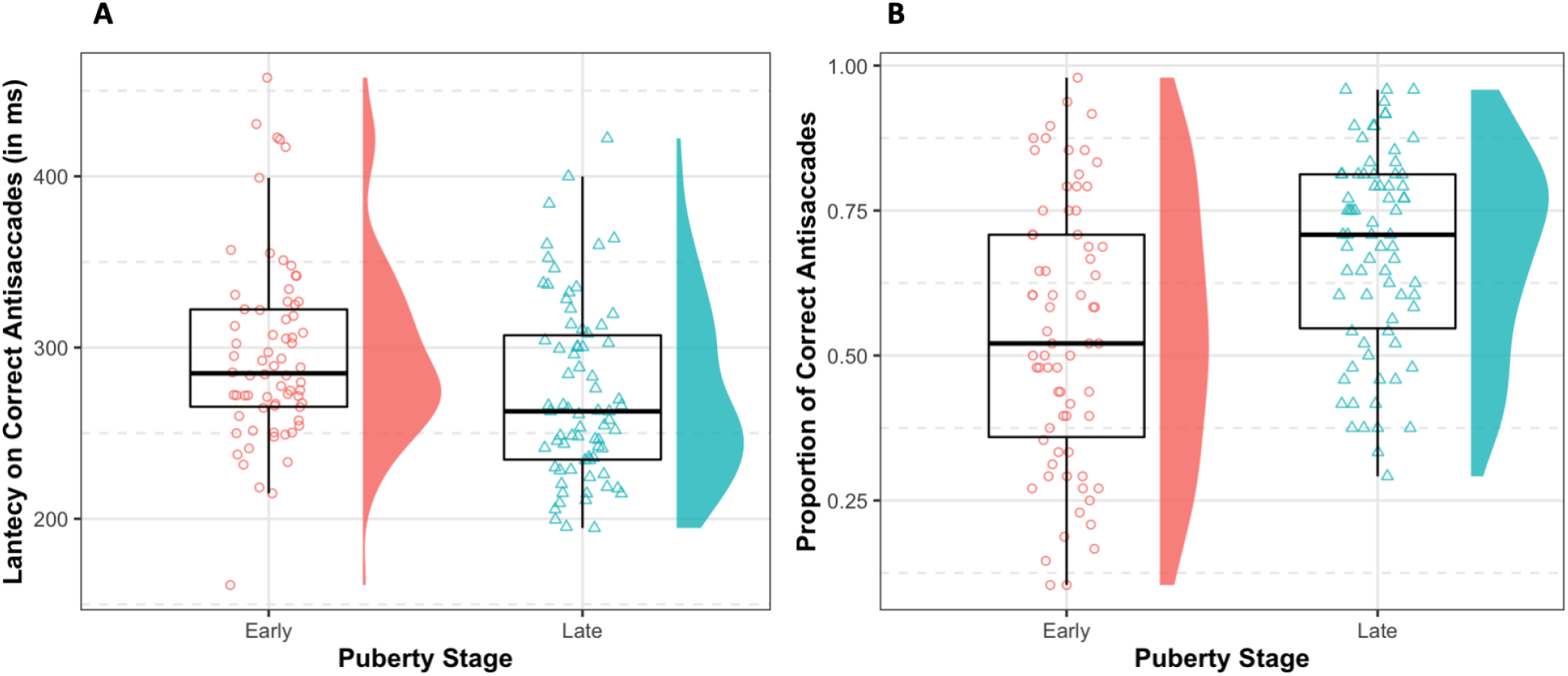
Distribution of antisaccade outcome variables separated by ‘Early’ puberty stage (transformed-PDS < 4) and ‘Late’ puberty stage (transformed-PDS ≥ 4) for (**A**) latency on correct trials and (**B**) proportion of correct trials.

**Figure S11.**
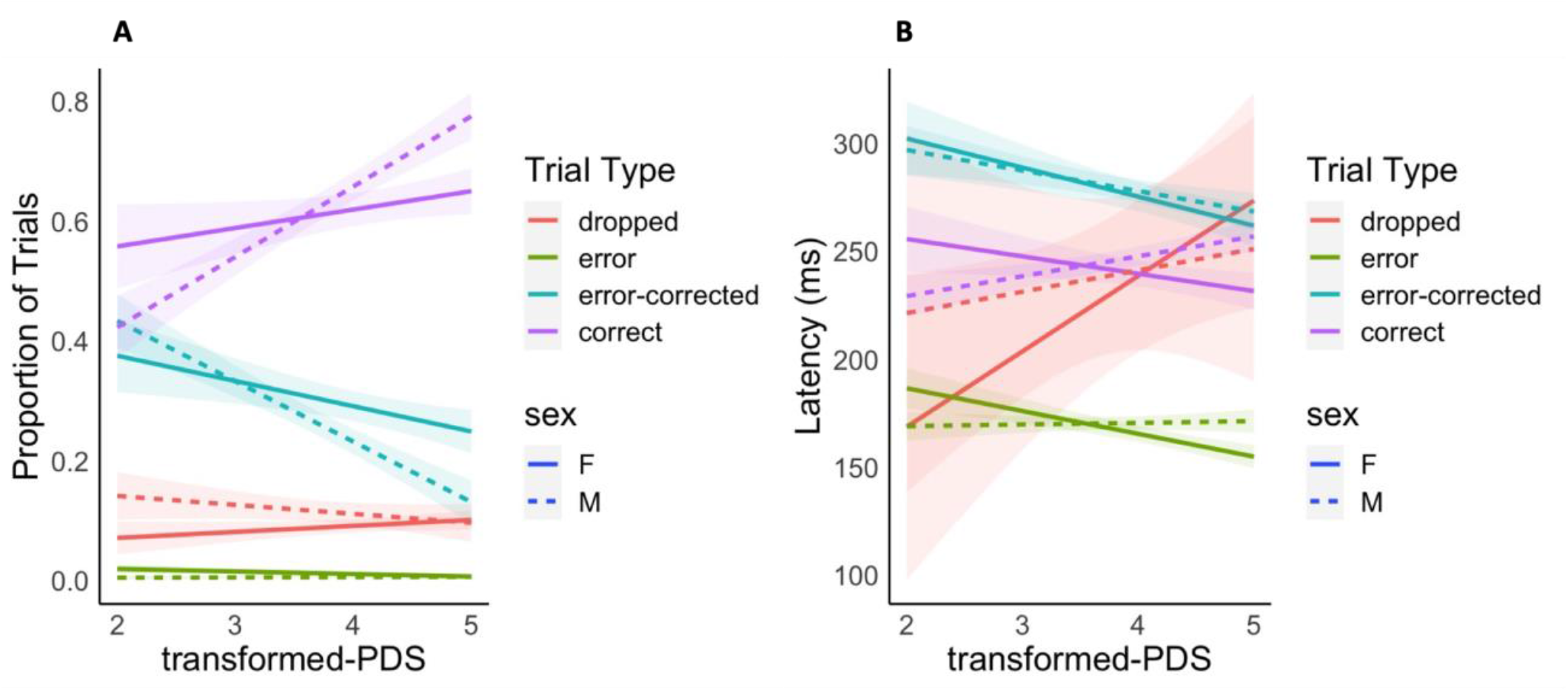
Plots depicting the relationship between pubertal maturation and the (**A**) proportion of antisaccades and (**B**) latency of antisaccades by trial type and sex.

**Figure S12.**
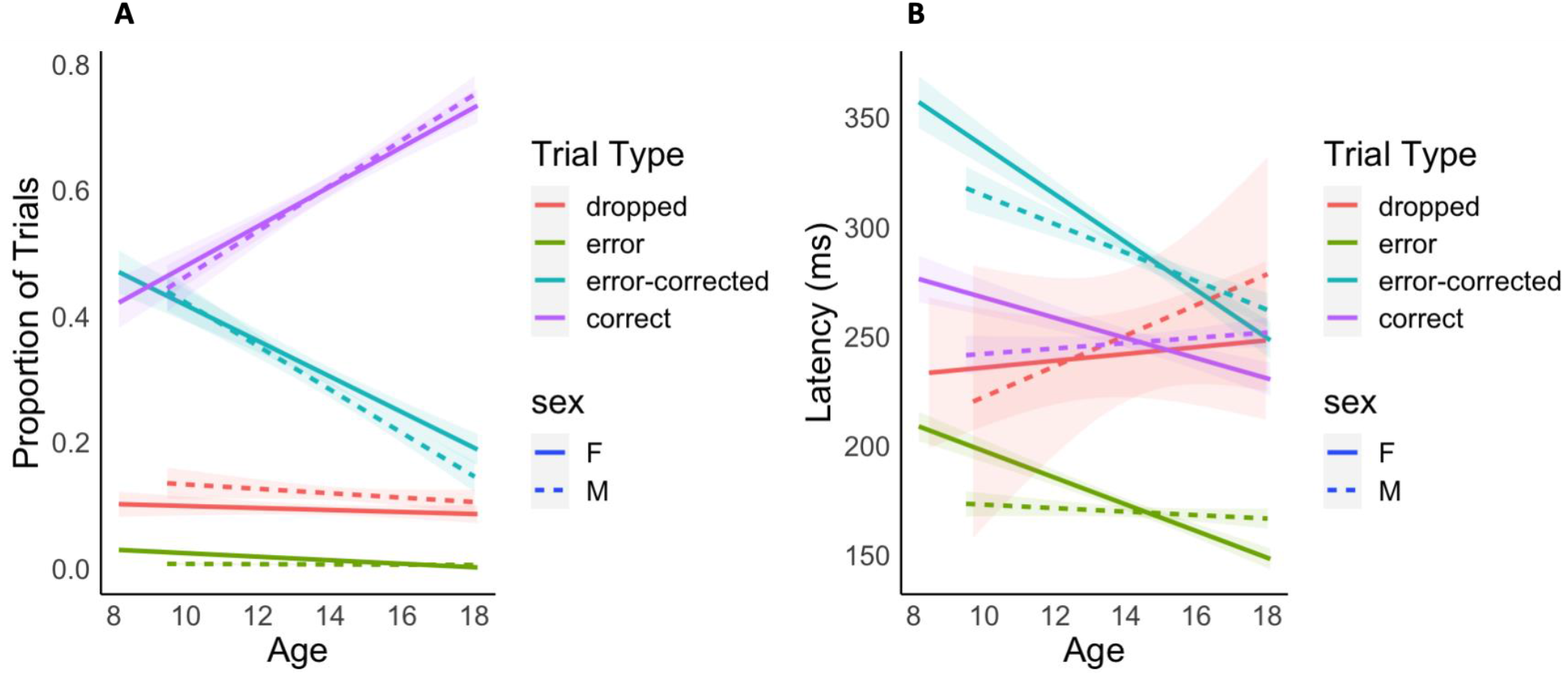
Plots depicting the relationship between age and the (**A**) proportion of antisaccades and (**B**) latency of antisaccades by trial type and sex.

## Supplemental Tables

**Table S1.** Sensitivity analyses testing the main effects of puberty controlling for percent of TRs censored in significant fronto-striatal connections.

| Striatal ROI | Frontal ROI | Puberty main effect |  | Puberty*sex interaction |  | Puberty (in males) |  | Puberty (in females) |  |
| --- | --- | --- | --- | --- | --- | --- | --- | --- | --- |
|  |  | F-statistic | *p*<sub>Bonferroni</sub> | F-statistic | *p*<sub>Bonferroni</sub> | F-statistic | *p*<sub>Bonferroni</sub> | F-statistic | *p*<sub>Bonferroni</sub> |
| NAcc |  |  |  |  |  |  |  |  |  |
|  | dIPFC | 16.90 | < .001*** |  |  |  |  |  |  |
|  | vlPFC | 14.59 | < .001*** |  |  |  |  |  |  |
| caudate |  |  |  |  |  |  |  |  |  |
|  | ant.vmPFC | 11.11 | < .001*** | 15.47 | < .001*** |  |  | 21.57 | < .001*** |
|  | rACC | 9.54 | < .001*** | 9.86 | < .001*** |  |  | 15.62 | < .001*** |
|  | vACC | 11.37 | < .001*** |  |  |  |  |  |  |
|  | dIPFC | 7.21 | .006** |  |  |  |  |  |  |
|  | vlPFC | 10.62 | < .001*** | 16.55 | < .001*** | 7.86 | < .001*** | 16.66 | < .001*** |
| putamen |  |  |  |  |  |  |  |  |  |
|  | vlPFC | 8.74 | .001** |  |  |  |  |  |  |

\* $p_{\text{Bonferroni}} < .05$ , \*\* $p_{\text{Bonferroni}} < .01$ , \*\*\* $p_{\text{Bonferroni}} < .001$

## References

Andersen, S.L., Thompson, A.P., Krenzel, E., Teicher, M.H., 2002. Pubertal changes in gonadal hormones do not underlie adolescent dopamine receptor overproduction. Psychoneuroendocrino 27, 683–691. https://doi.org/10.1016/s0306-4530(01)00069-5

Anderson, J.S., Ferguson, M.A., Lopez-Larson, M., Yurgelun-Todd, D., 2011. Reproducibility of Single-Subject Functional Connectivity Measurements. Am J Neuroradiol 32, 548–555. https://doi.org/10.3174/ajnr.a2330

Angold, A., Worthman, C.W., 1993. Puberty onset of gender differences in rates of depression: a developmental, epidemiologic and neuroendocrine perspective. J Affect Disorders 29, 145–158. https://doi.org/10.1016/0165-0327(93)90029-j

Bell, M.R., Meerts, S.H., Sisk, C.L., 2013. Adolescent brain maturation is necessary for adult-typical mesocorticolimbic responses to a rewarding social cue. Dev Neurobiol 73, 856–69. https://doi.org/10.1002/dneu.22106

Birn, R.M., Molloy, E.K., Patriat, R., Parker, T., Meier, T.B., Kirk, G.R., Nair, V.A., Meyerand, M.E., Prabhakaran, V., 2013. The effect of scan length on the reliability of resting-state fMRI connectivity estimates. Neuroimage 83, 550–558. https://doi.org/10.1016/j.neuroimage.2013.05.099

Blakemore, S.-J., 2008. The social brain in adolescence. Nat Rev Neurosci 9, 267–277. https://doi.org/10.1038/nrn2353

Bos, W. van den, Cohen, M.X., Kahnt, T., Crone, E.A., 2012. Striatum–Medial Prefrontal Cortex Connectivity Predicts Developmental Changes in Reinforcement Learning. Cereb Cortex 22, 1247–1255. https://doi.org/10.1093/cercor/bhr198

Braams, B.R., Duijvenvoorde, A.C.K. van, Peper, J.S., Crone, E.A., 2015. Longitudinal Changes in Adolescent Risk-Taking: A Comprehensive Study of Neural Responses to Rewards, Pubertal Development, and Risk-Taking Behavior. J Neurosci 35, 7226–7238. https://doi.org/10.1523/jneurosci.4764-14.2015

Calabro, F.J., Murty, V.P., Jalbrzikowski, M., Tervo-Clemmens, B., Luna, B., 2019. Development of Hippocampal-Prefrontal Cortex Interactions through Adolescence. Cereb Cortex New York N Y 1991 30, 1548–1558. https://doi.org/10.1093/cercor/bhz186

Cao, Z., Ottino-Gonzalez, J., Cupertino, R.B., Juliano, A., Chaarani, B., Banaschewski, T., Bokde, A.L.W., Quinlan, E.B., Desrivières, S., Flor, H., Grigis, A., Gowland, P., Heinz, A., Brühl, R., Martinot, J.-L., Martinot, M.-L.P., Artiges, E., Nees, F., Orfanos, D.P., Paus, T., Poustka, L., Hohmann, S., Millenet, S., Fröhner, J.H., Robinson, L., Smolka, M.N., Walter, H., Winterer, J., Schumann, G., Whelan, R., Mackey, S., Garavan, H., consortiu, I., 2021. Characterizing reward system neural trajectories from adolescence to young adulthood. Dev Cog Neuro 52, 101042. https://doi.org/10.1016/j.dcn.2021.101042

Cox, R.W., 1996. AFNI: software for analysis and visualization of functional magnetic resonance neuroimages. Comput Biomed Res Int J 29, 162–73. https://doi.org/10.1006/cbmr.1996.0014

Crone, E.A., 2014. The role of the medial frontal cortex in the development of cognitive and social-affective performance monitoring. Psychophysiology 51, 943–50. https://doi.org/10.1111/psyp.12252

Dalsgaard, S., Thorsteinsson, E., Trabjerg, B.B., Schullehner, J., Plana-Ripoll, O., Brikell, I., Wimberley, T., Thygesen, M., Madsen, K.B., Timmerman, A., Schendel, D., McGrath, J.J., Mortensen, P.B., Pedersen, C.B., 2020. Incidence Rates and Cumulative Incidences of the Full Spectrum of Diagnosed Mental Disorders in Childhood and Adolescence. Jama Psychiat 77, 155–164. https://doi.org/10.1001/jamapsychiatry.2019.3523

Delevich, K., Klinger, M., Okada, N.J., Wilbrecht, L., 2021. Coming of age in the frontal cortex: The role of puberty in cortical maturation. Semin Cell Dev Biol 118, 64–72. https://doi.org/10.1016/j.semcdb.2021.04.021

Dijk, K.R.A.V., Hedden, T., Venkataraman, A., Evans, K.C., Lazar, S.W., Buckner, R.L., 2010. Intrinsic Functional Connectivity As a Tool For Human Connectomics: Theory, Properties, and Optimization. J Neurophysiol 103, 297–321. https://doi.org/10.1152/jn.00783.2009

Drzewiecki, C.M., Willing, J., Juraska, J.M., 2016. Synaptic number changes in the medial prefrontal cortex across adolescence in male and female rats: A role for pubertal onset. Synapse 70, 361–368. https://doi.org/10.1002/syn.21909

Ernst, M., Nelson, E.E., Jazbec, S., McClure, E.B., Monk, C.S., Leibenluft, E., Blair, J., Pine, D.S., 2005. Amygdala and nucleus accumbens in responses to receipt and omission of gains in adults and adolescents. Neuroimage 25, 1279–91. https://doi.org/10.1016/j.neuroimage.2004.12.038

Fair, D.A., Schlaggar, B.L., Cohen, A.L., Miezin, F.M., Dosenbach, N.U.F., Wenger, K.K., Fox, M.D., Snyder, A.Z., Raichle, M.E., Petersen, S.E., 2007. A method for using blocked and event-related fMRI data to study “resting state” functional connectivity. Neuroimage 35, 396–405. https://doi.org/10.1016/j.neuroimage.2006.11.051

Fan, L., Li, H., Zhuo, J., Zhang, Y., Wang, J., Chen, L., Yang, Z., Chu, C., Xie, S., Laird, A.R., Fox, P.T., Eickhoff, S.B., Yu, C., Jiang, T., 2016. The Human Brainnetome Atlas: A New Brain Atlas Based on Connectional Architecture. Cereb Cortex New York N Y 1991 26, 3508–26. https://doi.org/10.1093/cercor/bhw157

Fareri, D.S., Gabard-Durnam, L., Goff, B., Flannery, J., Gee, D.G., Lumian, D.S., Caldera, C., Tottenham, N., 2015. Normative development of ventral striatal resting state connectivity in humans. Neuroimage 118, 422–437. https://doi.org/10.1016/j.neuroimage.2015.06.022

Fernandez–Galaz, M.C., Morschl, E., Chowen, J.A., Torres–Aleman, I., Naftolin, F., Garcia– Segura, L.M., 1997. Role of Astroglia and Insulin-Like Growth Factor-I in Gonadal Hormone-Dependent Synaptic Plasticity. Brain Res Bull 44, 525–531. https://doi.org/10.1016/s0361-9230(97)00238-4

Fortin, J.-P., Cullen, N., Sheline, Y.I., Taylor, W.D., Aselcioglu, I., Cook, P.A., Adams, P., Cooper, C., Fava, M., McGrath, P.J., McInnis, M., Phillips, M.L., Trivedi, M.H., Weissman, M.M., Shinohara, R.T., 2018. Harmonization of cortical thickness measurements across scanners and sites. Neuroimage 167, 104–120. https://doi.org/10.1016/j.neuroimage.2017.11.024

Galvan, A., Hare, T.A., Parra, C.E., Penn, J., Voss, H., Glover, G., Casey, B.J., 2006. Earlier development of the accumbens relative to orbitofrontal cortex might underlie risk-taking behavior in adolescents. J Neurosci Official J Soc Neurosci 26, 6885–92. https://doi.org/10.1523/jneurosci.1062-06.2006

Geier, C.F., Luna, B., 2012. Developmental Effects of Incentives on Response Inhibition: Motivated Response Inhibition. Child Dev 83, 1262–1274. https://doi.org/10.1111/j.1467-8624.2012.01771.x

Geier, C.F., Terwilliger, R., Teslovich, T., Velanova, K., Luna, B., 2009. Immaturities in Reward Processing and Its Influence on Inhibitory Control in Adolescence. Cereb Cortex 20, 1613–1629. https://doi.org/10.1093/cercor/bhp225

Glasser, M.F., Coalson, T.S., Robinson, E.C., Hacker, C.D., Harwell, J., Yacoub, E., Ugurbil, K., Andersson, J., Beckmann, C.F., Jenkinson, M., Smith, S.M., Essen, D.C.V., 2016. A multi-modal parcellation of human cerebral cortex. Nature 536, 171–178. https://doi.org/10.1038/nature18933

Goddings, A.-L., Mills, K.L., Clasen, L.S., Giedd, J.N., Viner, R.M., Blakemore, S.-J., 2014. The influence of puberty on subcortical brain development. Neuroimage 88, 242–251. https://doi.org/10.1016/j.neuroimage.2013.09.073

Grahn, J.A., Parkinson, J.A., Owen, A.M., 2008. The cognitive functions of the caudate nucleus. Prog Neurobiol 86, 141–155. https://doi.org/10.1016/j.pneurobio.2008.09.004

Hallquist, M.N., Hwang, K., Luna, B., 2013. The nuisance of nuisance regression: Spectral misspecification in a common approach to resting-state fMRI preprocessing reintroduces noise and obscures functional connectivity. Neuroimage 82, 208–225. https://doi.org/10.1016/j.neuroimage.2013.05.116

Hanlon, C.A., Dowdle, L.T., Austelle, C.W., DeVries, W., Mithoefer, O., Badran, B.W., George, M.S., 2015. What goes up, can come down: Novel brain stimulation paradigms may attenuate craving and craving-related neural circuitry in substance dependent individuals. Brain Res 1628, 199–209. https://doi.org/10.1016/j.brainres.2015.02.053

Harms, M.B., Casement, M.D., Teoh, J.Y., Ruiz, S., Scott, H., Wedan, R., Quevedo, K., 2019. Adolescent suicide attempts and ideation are linked to brain function during peer interactions. Psychiatry Res Neuroimaging 289, 1–9. https://doi.org/10.1016/j.pscychresns.2019.05.001

Harsay, H.A., Cohen, M.X., Oosterhof, N.N., Forstmann, B.U., Mars, R.B., Ridderinkhof, K.R., 2011. Functional Connectivity of the Striatum Links Motivation to Action Control in Humans. J Neurosci 31, 10701–10711. https://doi.org/10.1523/jneurosci.5415-10.2011

Hebbard, P.C., King, R.R., Malsbury, C.W., Harley, C.W., 2003. Two organizational effects of pubertal testosterone in male rats: transient social memory and a shift away from long-term potentiation following a tetanus in hippocampal CA1. Exp Neurol 182, 470–5. https://doi.org/10.1016/s0014-4886(03)00119-5

Ho, T.C., Colich, N.L., Sisk, L.M., Oskirko, K., Jo, B., Gotlib, I.H., 2020. Sex differences in the effects of gonadal hormones on white matter microstructure development in adolescence. Dev Cog Neuro 42, 100773. https://doi.org/10.1016/j.dcn.2020.100773

Ho, Tiffany C, Gifuni, A.J., Gotlib, I.H., 2021. Psychobiological risk factors for suicidal thoughts and behaviors in adolescence: a consideration of the role of puberty. Mol Psychiatr 27, 606–623. https://doi.org/10.1038/s41380-021-01171-5

Ho, Tiffany C., Teresi, G.I., Ojha, A., Walker, J.C., Kirshenbaum, J.S., Singh, M.K., Gotlib, I.H., 2021. Smaller caudate gray matter volume is associated with greater implicit suicidal ideation in depressed adolescents. J Affect Disorders 278, 650–657. https://doi.org/10.1016/j.jad.2020.09.046

Hu, S., Ide, J.S., Chao, H.H., Castagna, B., Fischer, K.A., Zhang, S., Li, C.R., 2018. Structural and functional cerebral bases of diminished inhibitory control during healthy aging. Hum Brain Mapp 39, 5085–5096. https://doi.org/10.1002/hbm.24347

Jahfari, S., Waldorp, L., Wildenberg, W.P.M. van den, Scholte, H.S., Ridderinkhof, K.R., Forstmann, B.U., 2011. Effective Connectivity Reveals Important Roles for Both the Hyperdirect (Fronto-Subthalamic) and the Indirect (Fronto-Striatal-Pallidal) Fronto-Basal Ganglia Pathways during Response Inhibition. J Neurosci 31, 6891–6899. https://doi.org/10.1523/jneurosci.5253-10.2011

Jones, R.M., Somerville, L.H., Li, J., Ruberry, E.J., Powers, A., Mehta, N., Dyke, J., Casey, B.J., 2014. Adolescent-specific patterns of behavior and neural activity during social reinforcement learning. Cognitive Affect Behav Neurosci 14, 683–697. https://doi.org/10.3758/s13415-014-0257-z

Kuhn, C., Johnson, M., Thomae, A., Luo, B., Simon, S.A., Zhou, G., Walker, Q.D., 2010. The emergence of gonadal hormone influences on dopaminergic function during puberty. Horm Behav 58, 122–137. https://doi.org/10.1016/j.yhbeh.2009.10.015

Ladouceur, C.D., 2012. Neural systems supporting cognitive-affective interactions in adolescence: the role of puberty and implications for affective disorders. Frontiers Integr Neurosci 6, 65. https://doi.org/10.3389/fnint.2012.00065

Ladouceur, C.D., Kerestes, R., Schlund, M.W., Shirtcliff, E.A., Lee, Y., Dahl, R.E., 2019. Neural systems underlying reward cue processing in early adolescence: The role of puberty and pubertal hormones. Psychoneuroendocrino 102, 281–291. https://doi.org/10.1016/j.psyneuen.2018.12.016

Ladouceur, C.D., Peper, J.S., Crone, E.A., Dahl, R.E., 2012. White matter development in adolescence: the influence of puberty and implications for affective disorders. Dev Cog Neuro 2, 36–54. https://doi.org/10.1016/j.dcn.2011.06.002

Larsen, B., Luna, B., 2018. Adolescence as a neurobiological critical period for the development of higher-order cognition. Neurosci Biobehav Rev 94, 179–195. https://doi.org/10.1016/j.neubiorev.2018.09.005

Larsen, B., Luna, B., 2014. In vivo evidence of neurophysiological maturation of the human adolescent striatum. Dev Cog Neuro 12, 74–85. https://doi.org/10.1016/j.dcn.2014.12.003

Luciana, M., Wahlstrom, D., Porter, J.N., Collins, P.F., 2012. Dopaminergic Modulation of Incentive Motivation in Adolescence: Age-Related Changes in Signaling, Individual Differences, and Implications for the Development of Self-Regulation. Dev Psychol 48, 844–861. https://doi.org/10.1037/a0027432

Luna, B., 2009. Developmental Changes in Cognitive Control through Adolescence. Adv Child Dev Behav 37, 233–278. https://doi.org/10.1016/s0065-2407(09)03706-9

Luna, B., Garver, K.E., Urban, T.A., Lazar, N.A., Sweeney, J.A., 2004. Maturation of Cognitive Processes From Late Childhood to Adulthood. Child Dev 75, 1357–1372. https://doi.org/10.1111/j.1467-8624.2004.00745.x

Luna, B., Thulborn, K.R., Munoz, D.P., Merriam, E.P., Garver, K.E., Minshew, N.J., Keshavan, M.S., Genovese, C.R., Eddy, W.F., Sweeney, J.A., 2001. Maturation of Widely Distributed Brain Function Subserves Cognitive Development. Neuroimage 13, 786–793. https://doi.org/10.1006/nimg.2000.0743

MacDonald, A.W., Cohen, J.D., Stenger, V.A., Carter, C.S., 2000. Dissociating the role of the dorsolateral prefrontal and anterior cingulate cortex in cognitive control. Sci New York N Y 288, 1835–8. https://doi.org/10.1126/science.288.5472.1835

Mackey, S., Petrides, M., 2014. Architecture and morphology of the human ventromedial prefrontal cortex. Eur J Neurosci 40, 2777–2796. https://doi.org/10.1111/ejn.12654

Mannella, F., Gurney, K., Baldassarre, G., 2013. The nucleus accumbens as a nexus between values and goals in goal-directed behavior: a review and a new hypothesis. Front Behav Neurosci 7, 135. https://doi.org/10.3389/fnbeh.2013.00135

Matsumoto, A., 1991. Synaptogenic action of sex steroids in developing and adult neuroendocrine brain. Psychoneuroendocrino 16, 25–40. https://doi.org/10.1016/0306-4530(91)90069-6

Mendle, J., Harden, K.P., Brooks-Gunn, J., Graber, J.A., 2010. Development’s tortoise and hare: Pubertal timing, pubertal tempo, and depressive symptoms in boys and girls. Dev Psychol 46, 1341–1353. https://doi.org/10.1037/a0020205

Morein-Zamir, S., Robbins, T.W., 2014. Fronto-striatal circuits in response-inhibition: Relevance to addiction. Brain Res 1628, 117–29. https://doi.org/10.1016/j.brainres.2014.09.012

Murty, V.P., Calabro, F., Luna, B., 2016. The role of experience in adolescent cognitive development: Integration of executive, memory, and mesolimbic systems. Neurosci Biobehav R 70, 46–58. https://doi.org/10.1016/j.neubiorev.2016.07.034

Murty, V.P., Shah, H., Montez, D., Foran, W., Calabro, F., Luna, B., 2018. Age-Related Trajectories of Functional Coupling between the VTA and Nucleus Accumbens Depend on Motivational State. J Neurosci 38, 7420–7427. <https://doi.org/10.1523/jneurosci.3508-17.2018>

Ochsner, K.N., Gross, J.J., 2008. Cognitive Emotion Regulation. Curr Dir Psychol Sci 17, 153–158. https://doi.org/10.1111/j.1467-8721.2008.00566.x

Oldehinkel, A.J., Verhulst, F.C., Ormel, J., 2010. Mental health problems during puberty: Tanner stage-related differences in specific symptoms. The TRAILS study. J Adolescence 34, 73–85. https://doi.org/10.1016/j.adolescence.2010.01.010

Ordaz, S.J., Foran, W., Velanova, K., Luna, B., 2013. Longitudinal Growth Curves of Brain Function Underlying Inhibitory Control through Adolescence. J Neurosci 33, 18109–18124. https://doi.org/10.1523/jneurosci.1741-13.2013

Ordaz, S.J., Fritz, B.L., Forbes, E.E., Luna, B., 2017. The influence of pubertal maturation on antisaccade performance. Developmental Sci 21, e12568. https://doi.org/10.1111/desc.12568

Padmanabhan, A., Geier, C.F., Ordaz, S.J., Teslovich, T., Luna, B., 2011. Developmental changes in brain function underlying the influence of reward processing on inhibitory control. Dev Cog Neuro 1, 517–529. https://doi.org/10.1016/j.dcn.2011.06.004

Parducz, A., Hajszan, T., MacLusky, N.J., Hoyk, Z., Csakvari, E., Kurunczi, A., Prange-Kiel, J., Leranth, C., 2006. Synaptic remodeling induced by gonadal hormones: Neuronal plasticity as a mediator of neuroendocrine and behavioral responses to steroids. Neuroscience 138, 977–985. https://doi.org/10.1016/j.neuroscience.2005.07.008

Parr, A.C., Calabro, F., Larsen, B., Tervo-Clemmens, B., Elliot, S., Foran, W., Olafsson, V., Luna, B., 2021. Dopamine-related striatal neurophysiology is associated with specialization of frontostriatal reward circuitry through adolescence. Prog Neurobiol 201, 101997. https://doi.org/10.1016/j.pneurobio.2021.101997

Parr, A.C., Calabro, F., Tervo-Clemmens, B., Larsen, B., Foran, W., Luna, B., 2022. Contributions of dopamine-related basal ganglia neurophysiology to the developmental effects of incentives on inhibitory control. Dev Cog Neuro 54, 101100. https://doi.org/10.1016/j.dcn.2022.101100

Patton, G.C., Viner, R., 2007. Pubertal transitions in health. Lancet Lond Engl 369, 1130–9. https://doi.org/10.1016/s0140-6736(07)60366-3

Paus, T., Keshavan, M., Giedd, J.N., 2008. Why do many psychiatric disorders emerge during adolescence? Nat Rev Neurosci 9, 947–57. https://doi.org/10.1038/nrn2513

Petersen, A.C., Crockett, L., Richards, M., Boxer, A., 1988. A self-report measure of pubertal status: Reliability, validity, and initial norms. J Youth Adolescence 17, 117–33. https://doi.org/10.1007/bf01537962

Pfeifer, J.H., Allen, N.B., 2021. Puberty Initiates Cascading Relationships Between Neurodevelopmental, Social, and Internalizing Processes Across Adolescence. Biol Psychiat 89, 99–108. https://doi.org/10.1016/j.biopsych.2020.09.002

Poon, J.A., Niehaus, C.E., Thompson, J.C., Chaplin, T.M., 2019. Adolescents’ pubertal development: Links between testosterone, estradiol, and neural reward processing. Horm Behav 114, 104504. https://doi.org/10.1016/j.yhbeh.2019.02.015

Primus, R.J., Kellogg, C.K., 1990. Gonadal hormones during puberty organize environment-related social interaction in the male rat. Horm Behav 24, 311–323. https://doi.org/10.1016/0018-506x(90)90012-m

Primus, R.J., Kellogg, C.K., 1989. Pubertal-related changes influence the development of environment-related social interaction in the male rat. Dev Psychobiol 22, 633–43. https://doi.org/10.1002/dev.420220608

Qu, Y., Galvan, A., Fuligni, A.J., Lieberman, M.D., Telzer, E.H., 2015. Longitudinal Changes in Prefrontal Cortex Activation Underlie Declines in Adolescent Risk Taking. J Neurosci Official J Soc Neurosci 35, 11308–14. https://doi.org/10.1523/jneurosci.1553-15.2015

Quach, A., Tervo-Clemmens, B., Foran, W., Calabro, F.J., Chung, T., Clark, D.B., Luna, B., 2020. Adolescent development of inhibitory control and substance use vulnerability: A longitudinal neuroimaging study. Dev Cog Neuro 42, 100771. https://doi.org/10.1016/j.dcn.2020.100771

Rasmussen, A.R., Wohlfahrt-Veje, C., Renzy-Martin, K.T. de, Hagen, C.P., Tinggaard, J., Mouritsen, A., Mieritz, M.G., Main, K.M., 2014. Validity of self-assessment of pubertal maturation. Pediatrics 135, 86–93. https://doi.org/10.1542/peds.2014-0793

Ravindranath, O., Ordaz, S.J., Padmanabhan, A., Foran, W., Jalbrzikowski, M., Calabro, F.J., Luna, B., 2020. Influences of affective context on amygdala functional connectivity during cognitive control from adolescence through adulthood. Dev Cog Neuro 45, 100836. https://doi.org/10.1016/j.dcn.2020.100836

Romeo, R.D., Sisk, C.L., 2001. Pubertal and seasonal plasticity in the amygdala. Brain Res 889, 71–7. https://doi.org/10.1016/s0006-8993(00)03111-5

Rubia, K., Smith, A.B., Woolley, J., Nosarti, C., Heyman, I., Taylor, E., Brammer, M., 2006. Progressive increase of frontostriatal brain activation from childhood to adulthood during event-related tasks of cognitive control. Hum Brain Mapp 27, 973–993. https://doi.org/10.1002/hbm.20237

Rudebeck, P.H., Rich, E.L., 2018. Orbitofrontal cortex. Curr Biol 28, R1083–R1088. https://doi.org/10.1016/j.cub.2018.07.018

Sato, S.M., Schulz, K.M., Sisk, C.L., Wood, R.I., 2008. Adolescents and androgens, receptors and rewards. Horm Behav 53, 647–658. https://doi.org/10.1016/j.yhbeh.2008.01.010

Schulz, K.M., Sisk, C.L., 2016. The organizing actions of adolescent gonadal steroid hormones on brain and behavioral development. Neurosci Biobehav Rev 70, 148–158. https://doi.org/10.1016/j.neubiorev.2016.07.036

Short, M.B., Rosenthal, S.L., 2008. Psychosocial Development and Puberty. Ann Ny Acad Sci 1135, 36–42. https://doi.org/10.1196/annals.1429.011

Shulman, E.P., Smith, A.R., Silva, K., Icenogle, G., Duell, N., Chein, J., Steinberg, L., 2016. The dual systems model: Review, reappraisal, and reaffirmation. Dev Cog Neuro 17, 103–117. https://doi.org/10.1016/j.dcn.2015.12.010

Siegel, J.S., Power, J.D., Dubis, J.W., Vogel, A.C., Church, J.A., Schlaggar, B.L., Petersen, S.E., 2013. Statistical improvements in functional magnetic resonance imaging analyses produced by censoring high-motion data points: Censoring High Motion Data in fMRI. Hum Brain Mapp 35, 1981–1996. https://doi.org/10.1002/hbm.22307

Silverman, M.H., Jedd, K., Luciana, M., 2015. Neural networks involved in adolescent reward processing: An activation likelihood estimation meta-analysis of functional neuroimaging studies. Neuroimage 122, 427–439. https://doi.org/10.1016/j.neuroimage.2015.07.083

Sisk, C.L., Foster, D.L., 2004. The neural basis of puberty and adolescence. Nat Neurosci 7, 1040–7. https://doi.org/10.1038/nn1326

Somerville, L.H., Jones, R.M., Ruberry, E.J., Dyke, J.P., Glover, G., Casey, B.J., 2012. The Medial Prefrontal Cortex and the Emergence of Self-Conscious Emotion in Adolescence. Psychol Sci 24, 1554–1562. https://doi.org/10.1177/0956797613475633

Spear, L.P., 2000. The adolescent brain and age-related behavioral manifestations. Neurosci Biobehav Rev 24, 417–463. https://doi.org/10.1016/s0149-7634(00)00014-2

Tervo-Clemmens, B., Quach, A., Luna, B., Foran, W., Chung, T., Bellis, M.D.D., Clark, D.B., 2017. Neural Correlates of Rewarded Response Inhibition in Youth at Risk for Problematic Alcohol Use. Front Behav Neurosci 11, 205. https://doi.org/10.3389/fnbeh.2017.00205

Thompson, T.L., Moss, R.L., 1994. Estrogen Regulation of Dopamine Release in the Nucleus Accumbens: Genomic- and Nongenomic-Mediated Effects. J Neurochem 62, 1750–1756. https://doi.org/10.1046/j.1471-4159.1994.62051750.x

van Duijvenvoorde, A.C.K., Achterberg, M., Braams, B.R., Peters, S., Crone, E.A., 2015. Testing a dual-systems model of adolescent brain development using resting-state connectivity analyses. Neuroimage 124, 409–20. https://doi.org/10.1016/j.neuroimage.2015.04.069

van Duijvenvoorde, A.C.K., Westhoff, B., Vos, F., Wierenga, L.M., Crone, E.A., 2019. A three-wave longitudinal study of subcortical–cortical restingstate connectivity in adolescence: Testing age- and puberty-related changes. Hum Brain Mapp 40, 3769–3783. https://doi.org/10.1002/hbm.24630

Vijayakumar, N., Whittle, S., Yücel, M., Dennison, M., Simmons, J., Allen, N.B., 2014. Thinning of the lateral prefrontal cortex during adolescence predicts emotion regulation in females. Soc Cogn Affect Neur 9, 1845–1854. https://doi.org/10.1093/scan/nst183

Voorn, P., Jorritsma-Byham, B., Dijk, C.V., Buijs, R.M., 1986. The dopaminergic innervation of the ventral striatum in the rat: A light- and electron-microscopical study with antibodies against dopamine. J Comp Neurology 251, 84–99. https://doi.org/10.1002/cne.902510106

Walhovd, K.B., Tamnes, C.K., Bjornerud, A., Due-Tonnessen, P., Holland, D., Dale, A.M., Fjell, A.M., 2014. Maturation of Cortico-Subcortical Structural Networks--Segregation and Overlap of Medial Temporal and Fronto-Striatal Systems in Development. Cereb Cortex 25, 1835–1841. https://doi.org/10.1093/cercor/bht424

Wood, S.N., 2013. A simple test for random effects in regression models. Biometrika 100, 1005–1010. https://doi.org/10.1093/biomet/ast038

Wood, S.N., Pya, N., Säfken, B., 2017. Smoothing Parameter and Model Selection for General Smooth Models. J Am Stat Assoc 111, 1548–1563. https://doi.org/10.1080/01621459.2016.1180986

Yuan, K., Yu, D., Cai, C., Feng, D., Li, Y., Bi, Y., Liu, J., Zhang, Y., Jin, C., Li, L., Qin, W., Tian, J., 2017. Frontostriatal circuits, resting state functional connectivity and cognitive control in internet gaming disorder. Addict Biol 22, 813–822. https://doi.org/10.1111/adb.12348

Zhang, F., Iwaki, S., 2020. Correspondence Between Effective Connections in the Stop-Signal Task and Microstructural Correlations. Front Hum Neurosci 14, 279. https://doi.org/10.3389/fnhum.2020.00279

Zhang, R., Geng, X., Lee, T.M.C., 2017. Large-scale functional neural network correlates of response inhibition: an fMRI meta-analysis. Brain Struct Funct 222, 3973–3990. https://doi.org/10.1007/s00429-017-1443-x

Zhou, Z., Chen, Y., Ding, M., Wright, P., Lu, Z., Liu, Y., 2009. Analyzing brain networks with PCA and conditional Granger causality. Hum Brain Mapp 30, 2197–2206. https://doi.org/10.1002/hbm.20661

